# Huntingtin Interacting Protein-1 expression is regulated via HIF2 axis in Lung Adenocarcinoma

**DOI:** 10.1101/2023.03.27.534427

**Authors:** Peeyush Prasad, Jonita Chongtham, Satyendra Chandra Tripathi, Nirmal Kumar Ganguly, Shivani Arora Mittal, Tapasya Srivastava

## Abstract

Non-Small Cell Lung Cancer (NSCLC) patients are diagnosed late when the disease has metastasized. *Kras* is a prevalent mutation in NSCLC besides EGFR and TP53 and targeted therapies against this have been challenging. We have explored deregulation of an endocytic adapter protein, Huntingtin Interacting Protein-1(HIP1) and its relevance in a *Kras* mutant lung adenocarcinoma cell line as a model system. HIP1 RNA expression is observed to be significantly reduced in high-grade and metastatic lung cancer patients as compared to low-grade tumours and this correlates with poor survival. HIP1 depletion followed by global proteome profiling in A549 cells identified metabolic pathways to be majorly upregulated, followed by RNA transport and surveillance, amongst others. HIP1 depletion also significantly increased anchorage independent growth and invasion of these cells. However, the EMT markers did not follow the canonical regulation. We observed E-Cadherin and Vimentin induction, which is suggestive of collective migration. Additionally, we observed a hypoxic microenvironment to induce HIP1 expression, mediated by Hypoxia Inducible Factor 2 (HIF2), suggesting that a HIF2-HIP1 axis can cause tumour suppression and needs further exploration.

## INTRODUCTION

Lung cancer is a leading cause of global cancer-related deaths, with a survival rate of less than 15% across all stages. Usually delayed diagnosis, when the disease has already metastasized, makes it difficult to target. Additionally, mutational status, clonal heterogeneity and diverse microenvironment conditions further complicates treatment. Studies also demonstrate that intratumoral hypoxia is clinically present in lung cancers, and adds another layer of complexity [1, 2]. *KRAS* is the most frequent mutation reported in non-small cell lung cancers (NSCLC). However, targeting KRAS has been challenging and targeted therapies for blocking functions of EGFR and ALK are being used.

Endocytosis, occurring in various forms, is important for cancer cell proliferation and metastasis. Endocytosis of ligand-receptor complex not only results in signalling attenuation, but can also result in its augmentation by persistence signalling inside endosomal vesicles and recycling of receptors to the membrane. Many of the endocytic proteins such as AP2, RhoA, Rac1 have validated roles as metastatic regulators [3]. NME1, MTSS1 and KISS1 are a few examples of metastatic suppressors which function by altering endocytosis [4]. Infact, targeting endocytosis in combination with Gefitinib to deregulate wtEGFR expression has shown optimistic results [5]. Due to their key roles in cancer progression and metastasis, it is important to elucidate the molecular functions of endocytic proteins in detail [6]. Huntingtin Interacting Protein 1(HIP1) is a key endocytic adapter protein involved in clathrin mediated endocytosis. It contains ANTH and Talin binding domains, thereby linking the endocytic and cytoskeletal processes, both critical for cancer cell progression. Oncogenic fusions of HIP1 with PDGFR and ALK in CML and NSCLC are reported [7, 8]. However, the molecular mechanism involving HIP1 in cell transformation remains undetermined. The prevailing theory is that HIP1 stabilises active Receptor Tyrosine Kinases during receptor-mediated endocytosis. HIP1 protein has also been shown to physically interact with EGFR, leading to its stabilisation and induction of its signalling [9]. Contrary to its oncogenic role, HIP1 is shown to have an anti-metastatic role in lung cancer. Only one study by Hsu et al. (2016) has explored HIP1 mediated regulation in lung cancer, where they found that it suppresses metastasis by repressing the Akt/beta catenin pathway [10]. Our study in an *in vitro* model system for lung adenocarcinoma not only corroborates their findings but also provides novel insights revealing HIP1 to regulate metabolism and anchorage independent growth. Also, we provide evidence for induction of HIP1 expression mediated by the HIF2 axis. These findings have implications for mechanistic regulation of HIF2 mediated tumour suppression of *Kras* mutant lung adenocarcinomas.

## Methods

### Study cohorts and patient survival data

To investigate the significance of HIP1 expression in lung cancer, gene expression data was obtained from the TCGA, GTeX and GEO datasets, which include 1865 primary, 8 metastatic tumours and 391 normal tissue samples [11]. Kaplan–Meier Plotter database was used to analyze the survival correlation between HIP1 mRNA expression and lung cancer patients. (Set parameters as median cutoff, hazard ratio (HR) with 95% confidence intervals (CIs), log rank P value and JetSet best probe with Affymetrix id 226364_at) [12].

### Cell Culture conditions

A549 cells were cultured at 37°C with 5% CO2 in 25 cm2 vented culture flasks in DMEM+ 10% Fetal Bovine Serum (FBS) media. Cells were passaged further and media changed after every 2- 3 days. Flasks were kept in the incubator (37 °C and 5 % CO2). Cells were cultured in both hypoxia ( 1% O2 concentration) and normoxia conditions.

### HIP1 Knockdown Assays

Cultured A549 cells were plated in sixLwell plates (2x10^5^ cells/well) one day prior to transfection so that they are around 80% confluent before transfection. Next day, the cells were transfected with HIP1 siRNAs using lipofectamine 3000, according to the manufacturer’s instructions. The plates were incubated at 37°C for 48 hrs.

Mutant stabilising constructs of HIF1 and HIF2 were commercially purchased and overexpressed by transient transfection using lipofectamine 3000, as per the manufacturer’s recommendations.

### Soft Agar Assay

Soft Agar Assay was performed by using 0.5% agar and 2% agar. First, 2% agar layer was formed in a 6-well plate and was allowed to solidify. After that, single cell suspension of cancer cells were made in 0.5% agar. This single cell suspension was poured on 2% solidified agar layer. Cells were allowed to proliferate for 21-28 days and were imaged by microscope at 10X.

### Immunoblotting

Validation of LC-MS data was done by western blot. Isolated protein lysate was separated on SDS-PAGE (8-12.5 % SDS-PAGE; 3h running at 100 V). Separated proteins on the gel were transferred to the PVDF membrane by applying current for 1.5 h in the transfer unit. Further, the membrane was incubated in the blocking solution (5% skim milk in PBS having 0.1% Tween-20 (1X PBST)) for 1 h. After three washing by 1X PBST, the membrane was incubated with the primary antibodies in 1X PBST overnight at 4 ^0^C. After overnight incubation, the membrane was washed with 1X PBST thrice and was incubated with secondary antibodies. Protein bands were visualised using ECL solution. Intensity of the bands was measured by ImageJ software [13].

### Ki67 staining

For performing Ki67 staining, knockdown cells were harvested after 48 h. Harvested cells were vortexed and while vortexing cold ethanol was added dropwise. After this, cells were incubated at -20°C for at least 2 h. After 2 h, cells were washed twice with a staining buffer by centrifugation. Cells were re-suspended again in a staining buffer and Ki67 antibody was added and incubated for 20-30 min in dark. After incubation, cells were washed with a staining buffer. Finally, supernatant was aspirated and 0.5 ml of staining buffer was added and processed for flow cytometry analysis.

### Matrigel Invasion Assay

Invasion assay was performed by using matrigel and culture insert. Cells were seeded in a 6-well plate. Transfection was carried out after 24 h at 80% confluency for 48 h. Next day, matrigel was thawed onto the ice overnight. After thawing, matrigel was coated onto the culture insert. Cultur insert plate was kept in an incubator (5% CO2 and 37 degree-centigrade) for 4-6 h. Transfected cells were harvested after 24 h and seeded onto the prepared matrigel coated culture insert. After this, the culture insert plate was kept in the incubator. After 24 hr of seeding on the coated culture insert, inserts were processed for imaging. For imaging cells containing culture inserts fixed by formaldehyde and permeabilized by methanol. Crystal violet solution was used for staining purposes. After cleaning the inner side of the insert’s well with a cotton swab, cells were imaged on a microscope at 10X by keeping the insert on a slide.

### MTT Assay

For the MTT assay, knockdown cells were seeded in a 96-well plate. After 24 h and 48 h of seeding, media was aspirated and MTT solution was added. Plate was incubated at 37°C for 3 hours. After 3 h, DMSO (MTT solvent) was added into each well and the plate was shaken for 15 min. at an orbital shaker. Finally, absorbance was read at OD=590 nm.

% viability was calculated by following formula: 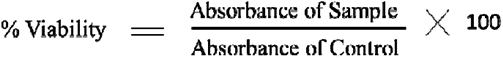

### Cell cycle Analysis

Knockdown cells were harvested and washed by 1X PBS. Cells were fixed in cold ethanol by adding ethanol drop wise while vortexing. Cells were further kept at -20^0^ C for 2 h. Cells were washed 2 times by 1X PBS by centrifugation. Cells were treated with RNase. 200 ul of Propidium Iodide (from 50 µg/ml stock solution) was added and flow cytometry analysis was performed by measuring forward scatter and side scatter to identify single cells.

### Trans-well migration assay

Cells were seeded in 6 well plates at a density approximately ∼2×10^5^ cells per well. After 24 h, cells were treated with siHIP1 and siScrambled for 48 hr. Transfected cells were seeded onto the upper chamber of culture insert and in the lower chamber 10% FBS media was added as a chemoattractant. Cells were allowed to migrate through the pores of the insert for 24 h. Finally, cells were fixed by 3.7% formaldehyde for 10 min, permeabilized by 100% methanol for 15 min and stained by 0.2% crystal violet for 10 minutes. Non-invaded cells were removed using a cotton swab from the inner chamber of the culture inserts. Invaded cells were imaged by using a microscope at 10X.

#### RNA isolation & qPCR

RNA was extracted according to the manufacturer’s protocol using Trizol reagent. Reverse transcription was performed by using a high capacity cDNA Reverse Transcription kit (Applied Biosystems). The mRNA was mixed with reverse transcriptase enzyme and incubated in Thermal Cycler (procedure: 25°C for 10 min, 37°C for 2h and 85°C for 5 s). For Real-Time PCR, a serially diluted standard curve and all cDNAs samples were amplified by using SYBR Green in Thermal Cycler (procedure: 95°C for 10 min, 35 cycles of 95°C for 15 s and 60°C for 1 min). Primers for housekeeping genes were used for internal control normalisation. A dissociation curve was included for checking the purity of PCR products.

### LC-MS (Proteomics)

2 x 10^5^ A549 cells were seeded in a 6-well plate. After 24 h, at 70-80% confluency cells were transfected by 20 nm siHIP1 and siScrambled using Lipofectamine 3000. After 48 h, cells were harvested and lysed. Protein lysate was quantified by BCA. 25 ug protein from each sample was reduced with 5 mM TCEP and further alkylated with 50 mM iodoacetamide and then digested with Trypsin (1:50, Trypsin/lysate ratio) for 16 h at 37 °C. Digests were cleaned using a C18 silica cartridge to remove the salt and dried using a speed vac. The dried pellet was resuspended in buffer A (5% acetonitrile, 0.1% formic acid). Experiments were performed on an Ultimate 3000 RSLCnano system coupled with a Thermo QE Plus. 1ug was loaded on a C18 column 50 cm, 3.0μm Easy-spray column (Thermo Fisher Scientific). Peptides were eluted with a 0–40% gradient of buffer B (80% acetonitrile, 0.1% formic acid) at a flow rate of 300 nl/min ) and injected for MS analysis. LC gradients were run for 100 minutes. MS1 spectra were acquired in the Orbitrap at 70k resolution. Dynamic exclusion was employed for 10 s excluding all charge states for a given precursor. MS2 spectra were acquired at 17500 resolutions. All samples were processed and RAW files generated were analyzed with Proteome Discoverer (v2.2) against the Uniprot Human reference proteome database. For Sequest and Amanda search, the precursor. and fragment mass tolerances were set at 10 ppm and 0.5 Da, respectively. The protease used to generate peptides, i.e. enzyme specificity was set for trypsin/P (cleavage at the C terminus of “K/R: unless followed by “P”) along with maximum missed cleavage value of two. Carbamidomethyl on cysteine as fixed modification and oxidation of methionine and N-terminal acetylation were considered as variable modifications for database search. Both peptide spectrum match and protein false discovery rate were set to 0.01 FDR.

### Bioinformatics analysis

Dysregulated proteins identified in the high-throughput experiment were analyzed by bioinformatics analysis. Gene Ontology (Biological process and KEGG pathways), was carried out by using the STRING tool [14]. Network analysis was done using Ingenuity Pathway Analysis (IPA).

## Results

### 1. Reduced HIP1 expression correlates with poor survival in advanced Lung Adenocarcinoma patients

A pan-cancer analysis from the TCGA database revealed that lung cancer exhibits significantly reduced levels of HIP1 in tumour tissue compared to normal tissues (Supl Fig.1). To further analyse the status of HIP1 expression within lung cancer, we compared HIP1 expression within subtypes as well as between normal, tumour and metastatic tissues. Results showed significantly decreased levels of HIP1 transcripts in lung cancer compared to normal tissues (Fig. 1A and 1B; Supl Fig 1). Simultaneously, comparison of normal, tumour, and metastatic tissues further revealed that HIP1 is downregulated in metastatic cases as compared to tumour tissues (Fig. 1C).

**Figure 1.**
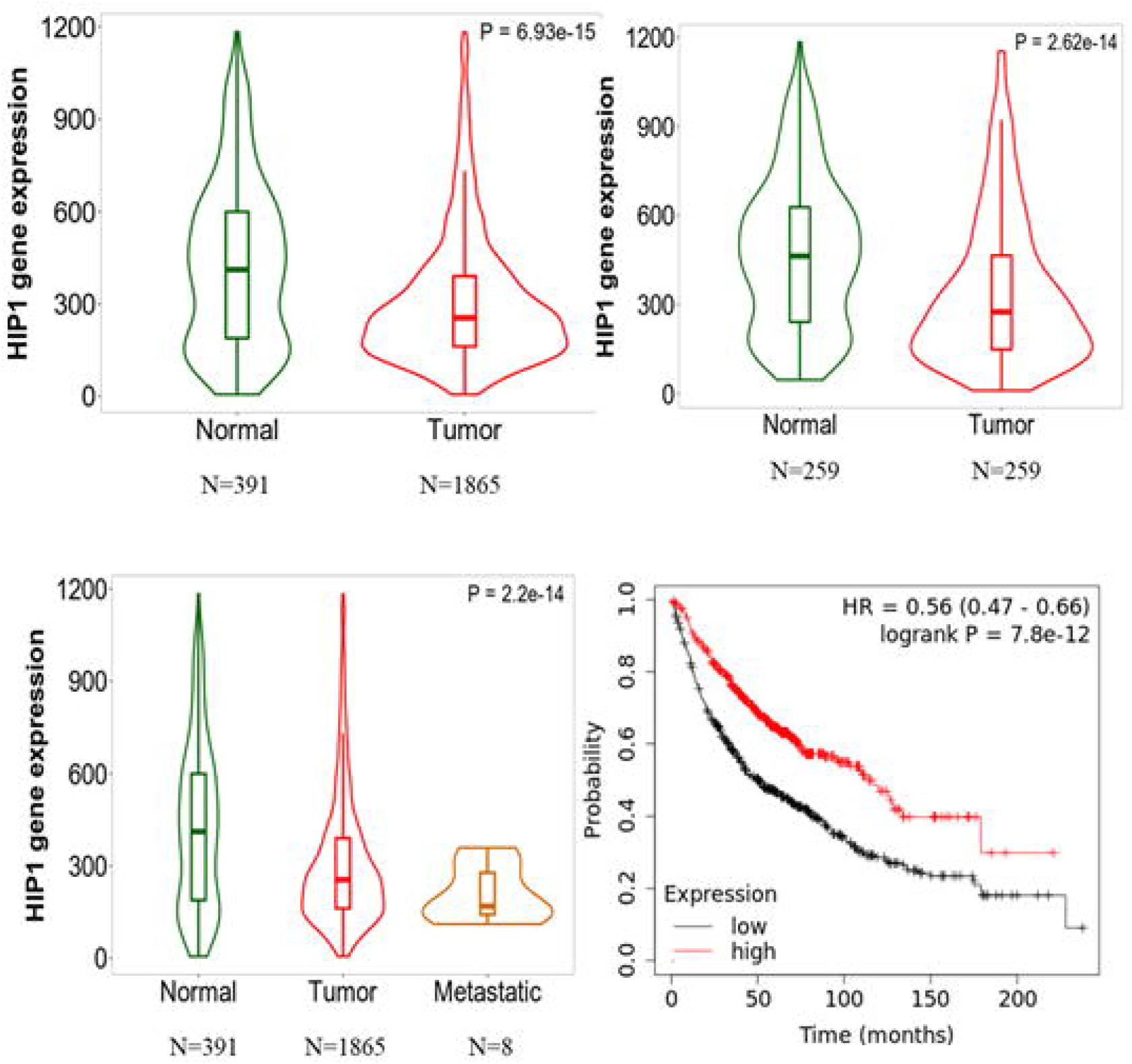
HIP1 expression in lung cancer: Violin plots of HIP 1 gene expression in Lung cancer (A) unpaired normal vs tumor (B) paired normal and tumor and (C) unpaired normal, tumor and metastatic gene array data. For unpaired samples Mann-Whitney U test and for paired samples, a paired Wilcoxon statistical test was used. (D) Kaplan-Meier survival curves of lung cancer patients with decreased expression of HIP 1 showed poor overall survival compared to those with increased HIP1 expression.

Kaplan-Meier survival analysis for 1144 lung cancer patients revealed that decreased HIP1 expression is significantly associated with poor prognosis (Fig.1D and Supl Fig 2). A median overall and progression free survival of 50 and 18 months was observed in lung cancer patients with low expression of HIP1 as compared to 112 and 45 months in the patients with high HIP1 expression, respectively.

### 2. HIP1 silencing identifies globally deregulated pathways

To understand the functional role of HIP1 in lung adenocarcinoma, we transiently silenced HIP1 expression in lung cancer cells followed by a global label-free quantitative proteomics. A total of 2,539 proteins were identified. Out of these, 263 were found upregulated and 139 proteins were found downregulated, using Log_2_ 2 fold change as cutoff. This data was validated by immunoblotting of two key down regulated proteins from the top 20 list, Succinate Dehydrogenase B (4 fold) and Prefoldin (2.5 fold) [Fig 2A]. Gene Ontology (GO) analysis was performed for the differentially regulated proteins using STRING database [13]. The majority of upregulated proteins were involved in metabolic pathways (18.83%). Others were found implicated in pathways such as RNA transport, Huntington’s disease, mRNA surveillance, endocytosis, mTOR signalling [Fig 2B]. The key down regulated biological processes were identified as protein localization, protein transport and cytoskeleton organisation, amongst others. Two downregulated KEGG pathways found were those of spliceosome and RNA transport (Supl. Fig 3 & Supl. Fig 4).

**Fig 2A.**
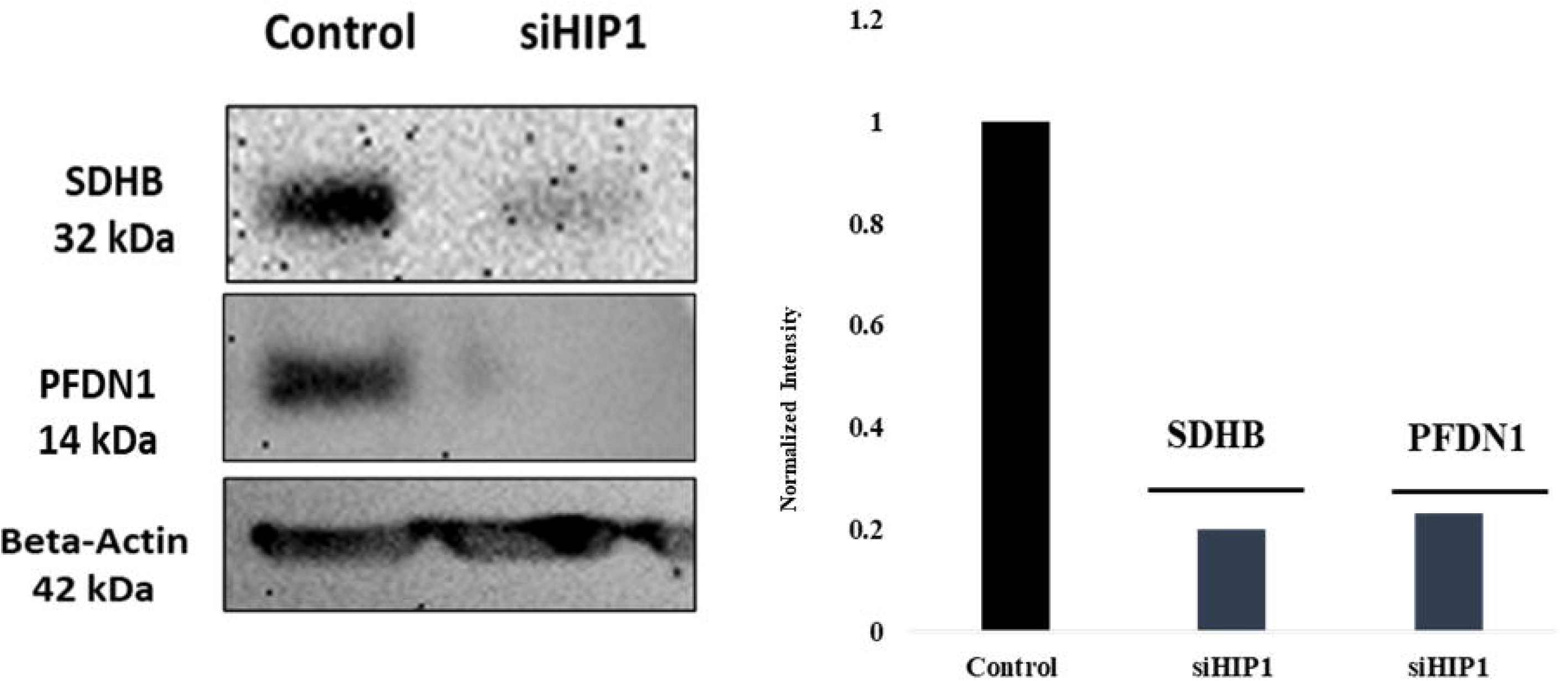
HIP1 siRNA (siHIPl)/scrambled siRNA (Control) was transfected in A549 cells for transient silencing. After 48 hrs HIP1 silenced cells were harvested and iimnunoblotted for HIP1, SDHB. PFDN1. SDHB and PFDN1 were both found downregulated in HIP1 silenced A549 cells.

**Figure 2B.**
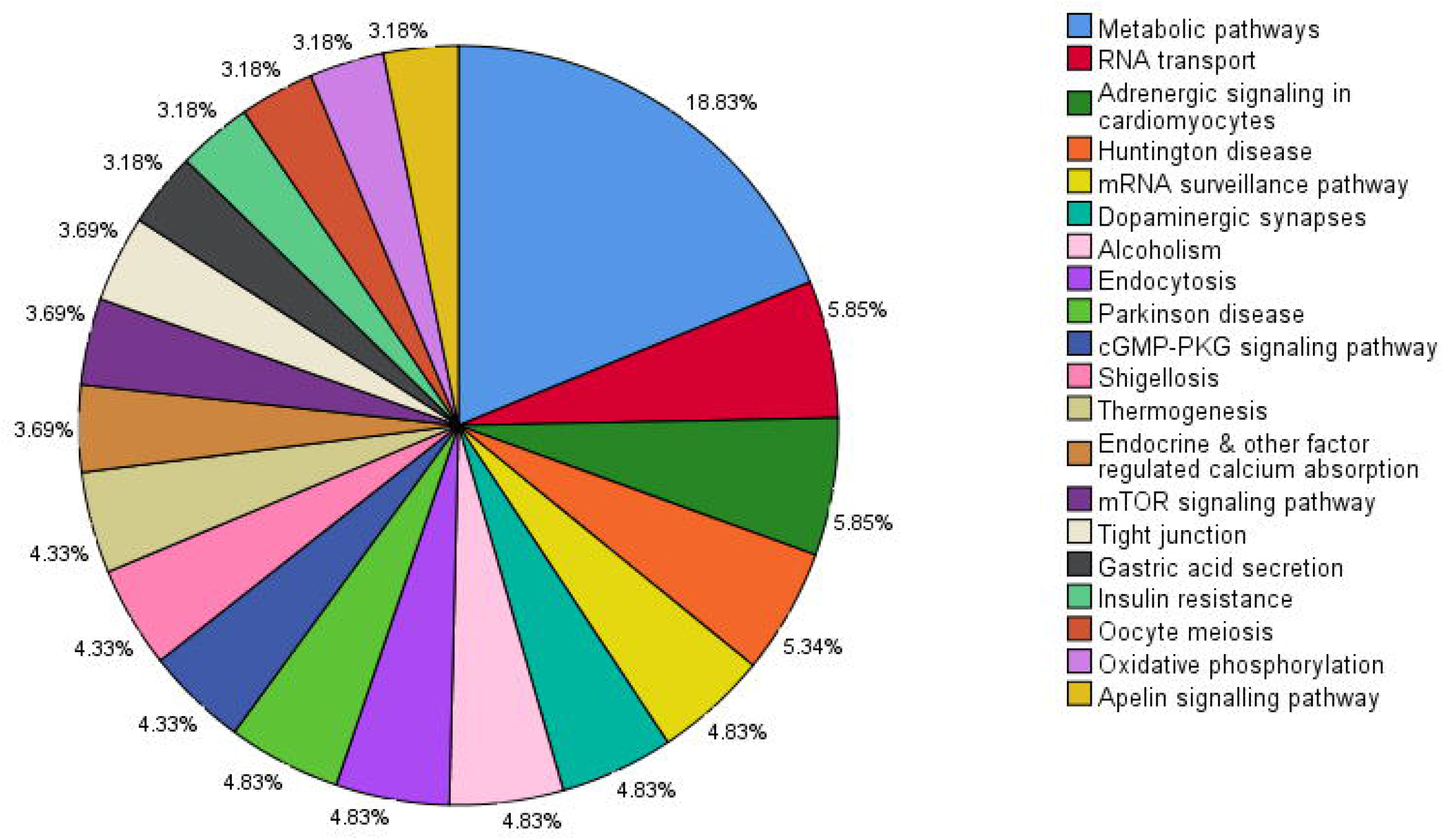
**Gene Ontology analysis of upregulated proteins after HIP1 knockown in A549 cells.** Majority of the upregulated proteins were found to be involved in various pathways such as metabolic (18.83%), RNA transport (5.85%), Huntington’s disease (5.34%) and Dopaminergic synapse (4.83%). Pie chart represents the top 20 KEGG pathways associated with upregulated proteins.

Using the Ingenuity Pathway Analysis (IPA) tool, the deregulated proteins were mostly associated with cellular assembly and organisation, metabolic disease, cellular movement and carbohydrate metabolism [Supplementary excel file 1]. Further, IPA identified activation of key regulators like NFE2L2, ESR2 and MYC and inhibition of regulators like miR-145-5p, miR-517- 5p, miR-497-3p and CDKN1B after HIP1 knockdown [Supplementary excel file 2]. The analysis revealed (Fig 2C), MYC induction as a key upstream activator, further activating NFE2L2 (NRF2), NFE2L1, MAFK, leading to phenotypic effects of cell proliferation, invasion and cell cycle progression (Fig 2D). Few other important activated components of this network are EGFR, NMAFK and cytokines such as IL15, IL4. Also, predicted mTORC1 signalling activation leads to induction of metabolic regulators like Phosphofructokinase, Lactate Dehydrogenase A and various Cyclin Dependent Kinases (CDK2/4/6), which could lead to increased survival and proliferation of cancer cells (Fig 2E).

**Figure 2C:**
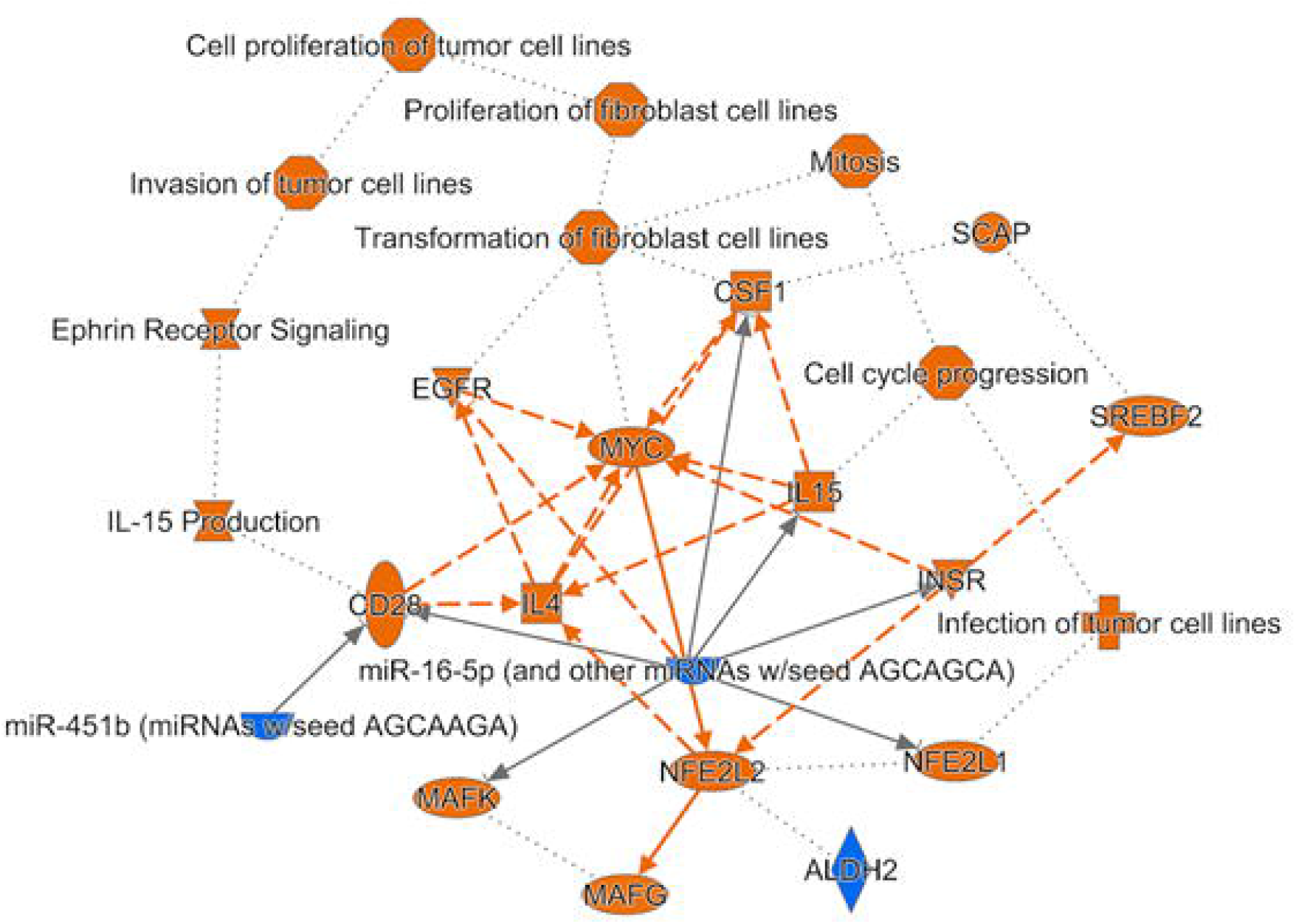
Graphical Summary of the upstream regulators after HIP1 knockdown. Differentially regulated proteins analysed using IPA identified various key regulators to be induced leading to net effect of increased cell proliferation, cell cycle progression and invasion of tumor cells.

**Figure 2D.**
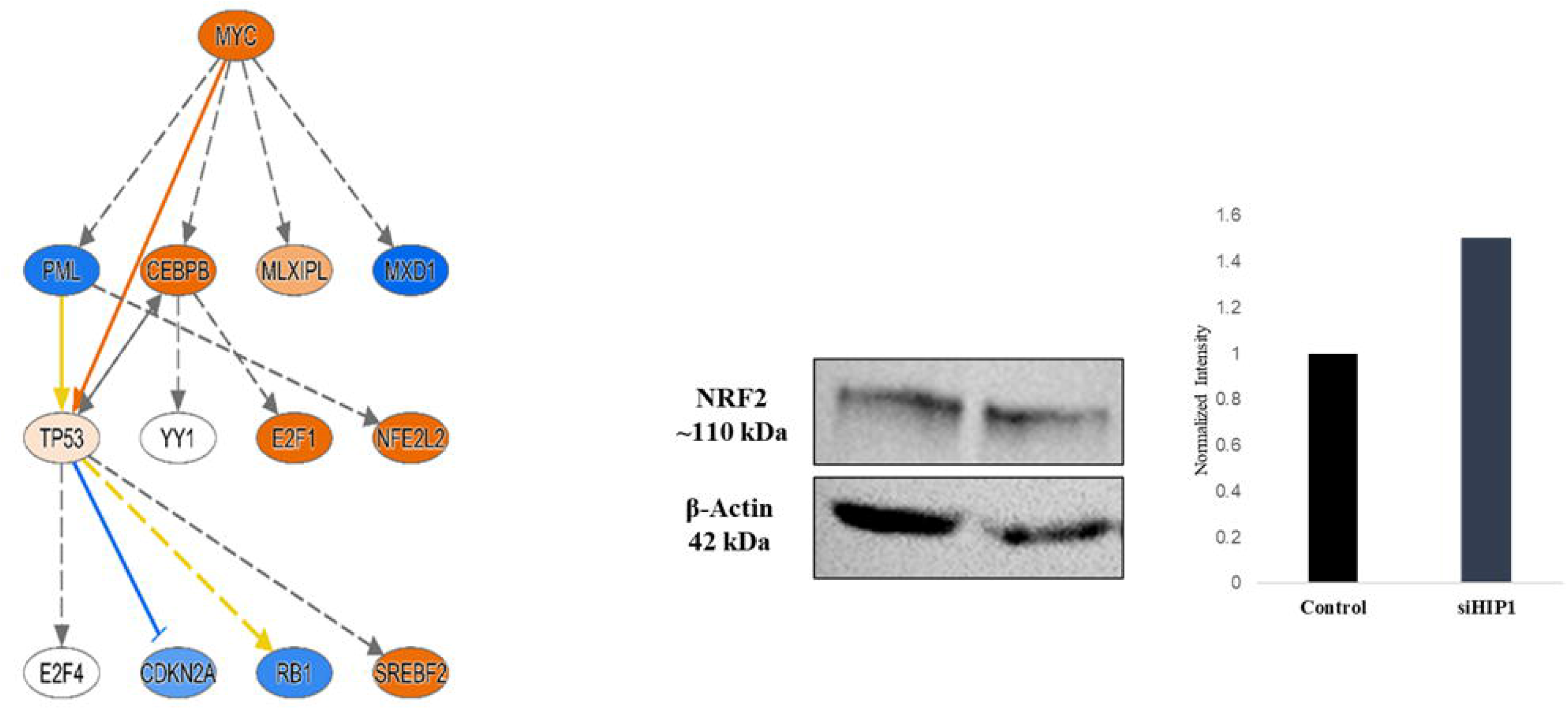
MYC is predicted as a key upstream regulator of differentially regulated proteins and is associated with proliferation, invasion, suivival and antioxidant response in HIP1 knockdown A549 cells. After HIP 1 downregulation. NRF2 (NFE2L2) is found upregulated by 1.5 fold by immunoblotting.

**Figure 2E.**
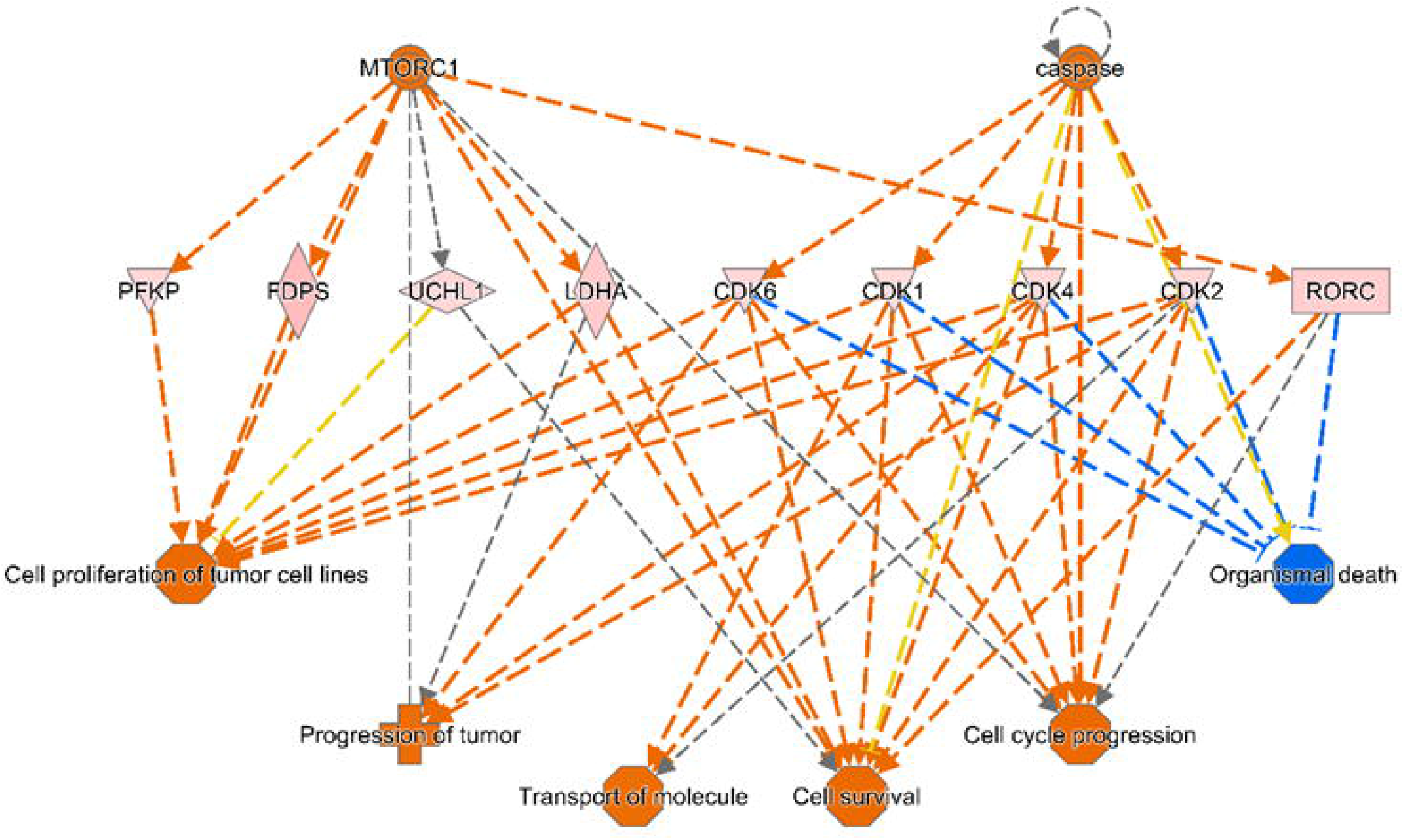
mTOR and caspase activation after HIP1 knockdown. Activation of mTORC and caspase signaling pathway leads to activation of proteins like PFKP, PDFS and CDK6 and RORC that are involved in cell proliferation, tumor progression and cell survival.

### 3. HIP1 silencing augments invasion and anchorage independent growth

Using insights from IPA analysis, phenotypic assays were performed to further confirm the role of HIP1 in lung cancer progression. HIP1 silencing induced the migratory and invasive properties of A549 cells. Transwell migration of HIP1 knockdown cells was significantly increased (1.5 fold). Also, in Matrigel invasion assay, we found higher (2 fold) invasive abilities of these cells (Fig 3A and 3B). Thus, reduced HIP1 expression in lung adenocarcinoma cells observed at higher stages induces their migratory and invasive abilities.

**Figure 3A.**
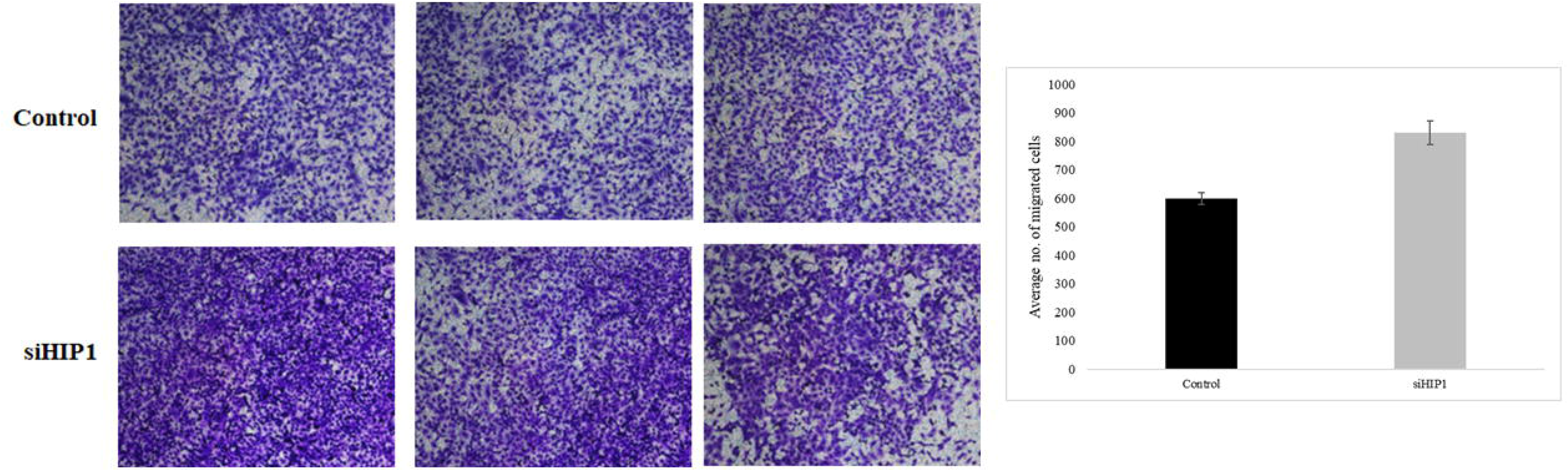
HIP1 siRNA (siHIPl)/scrambled siRNA (Control) was transfected in A549 cells. After 48 hrs HIP1 silenced cells were seeded onto culture inserts. These cells were then fixed, penneabilized and stained with crystal violet (blue color) to look at the migrated cells. Average number of migrated cells are plotted as a bar diagram (P=0.07)

**Figure 3B.**
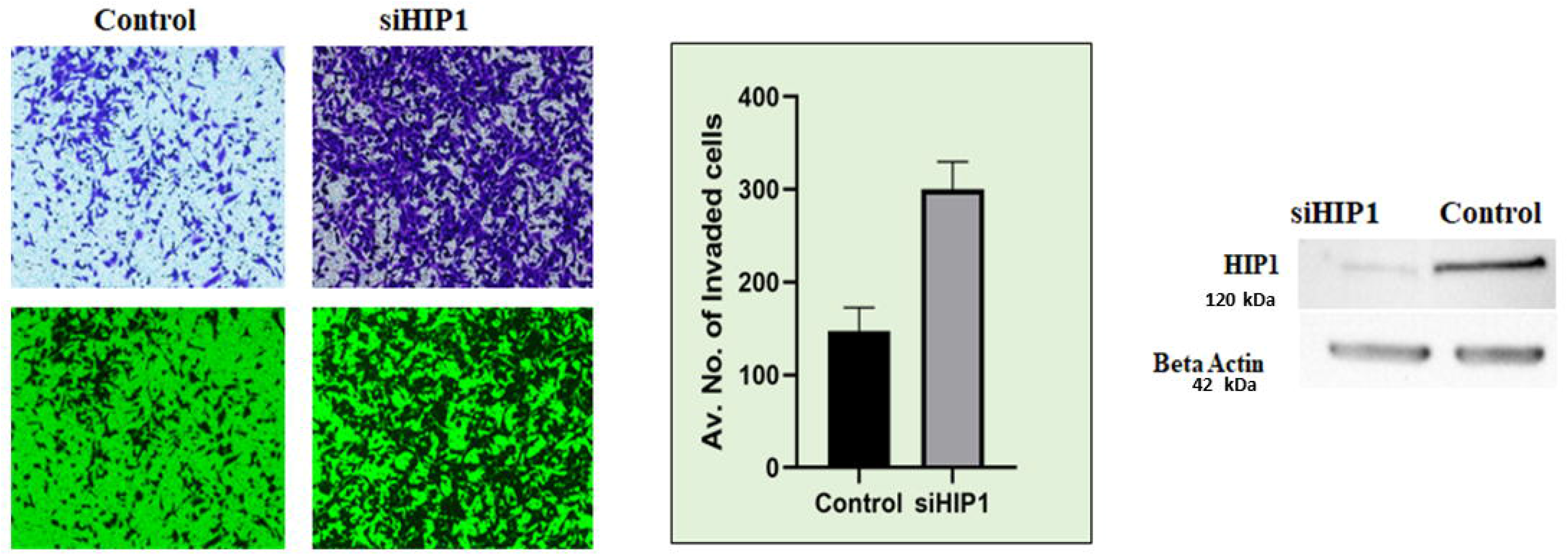
HIP1 siRNA (siHIPl )/scrambled siRNA (Control) along with siGLOW (green) was transfected in A549 cells. Post 48 hrs, HIP1 silenced cells were seeded onto matrigel coated culture inserts. These cells were then fixed, stained with crystal violet (blue color) after twenty four hours to look at the invaded cells. Also, HIP1 silencing in the same cells was confirmed using immunoblotting. Average number of invaded cells are plotted as a bar diagram (P=0.05).

Also, colony forming ability of HIP1 knockdown cells was assessed by culturing single cells in anchorage dependent and anchorage independent conditions. We observed significantly larger colony sizes in anchorage independent conditions (Fig 3D and 3E). On the contrary, HIP1 knockdown cells exhibited significantly reduced mitochondrial redox potential as implied by reduced MTT readings (Fig 3F). Also, no change in the cell cycle stages or Ki67 protein levels was observed (Supl Fig 6 and Supl Fig 7). This implies that HIP1 depletion in lung cancer activates alternate pathways of anchorage independent survival and growth.

**Figure 3C.**
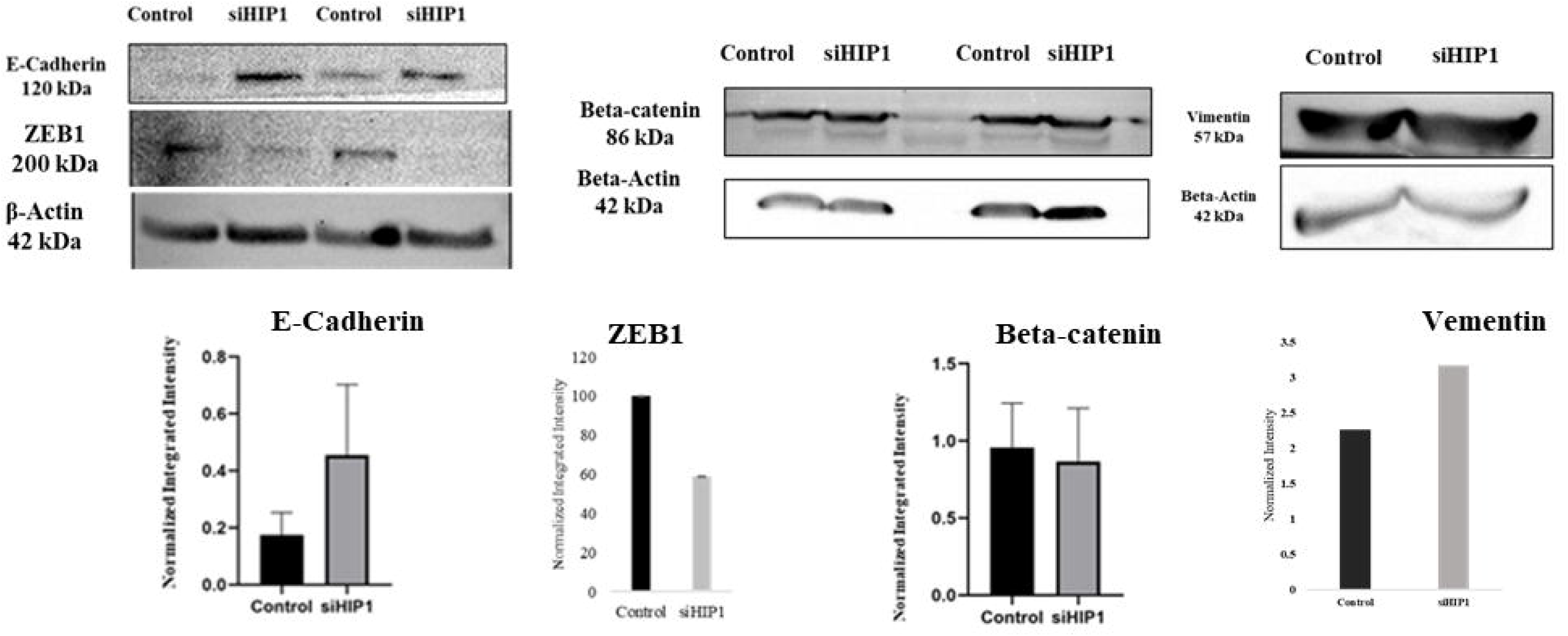
A549 cells were transfected with scrambled siRNA (control) and HIP1 siRNA (siHIPl). After confirming HIP1 downregulation, levels of E-cad, Beta-catenin, vimentin and ZEB1 was confirmed using immunoblot. No change in expression of beta catenin. increased expression of E-cad and vimentin and decreased expression of ZEB1 was found upon downregulation of HIP 1.

**Figure 3D.**
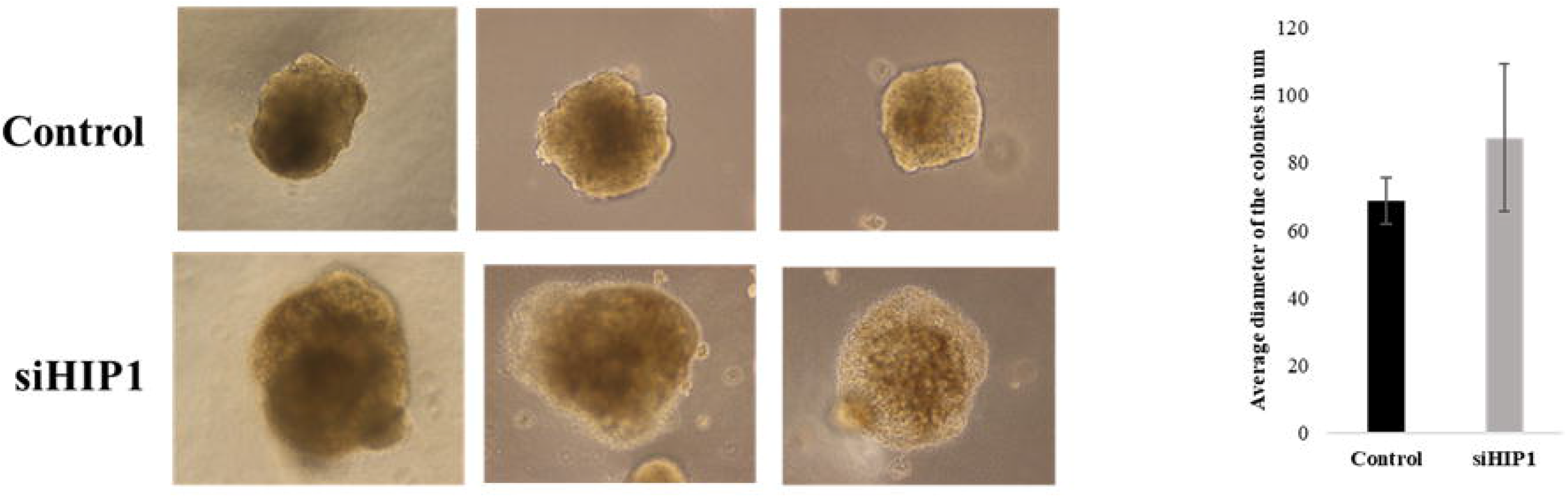
HIP1 siRNA (siHIPl)/scrambled siRNA(Control) was transfected in A549 cells. After 48 Ins cells were resuspended in 0.5% agar and poured onto the 2% solid agar surface. Cells were allowed to make colonies for 21 days and average diameter of colonies are plotted as a bar diagram (P=0.113).

**Figure 3E.**
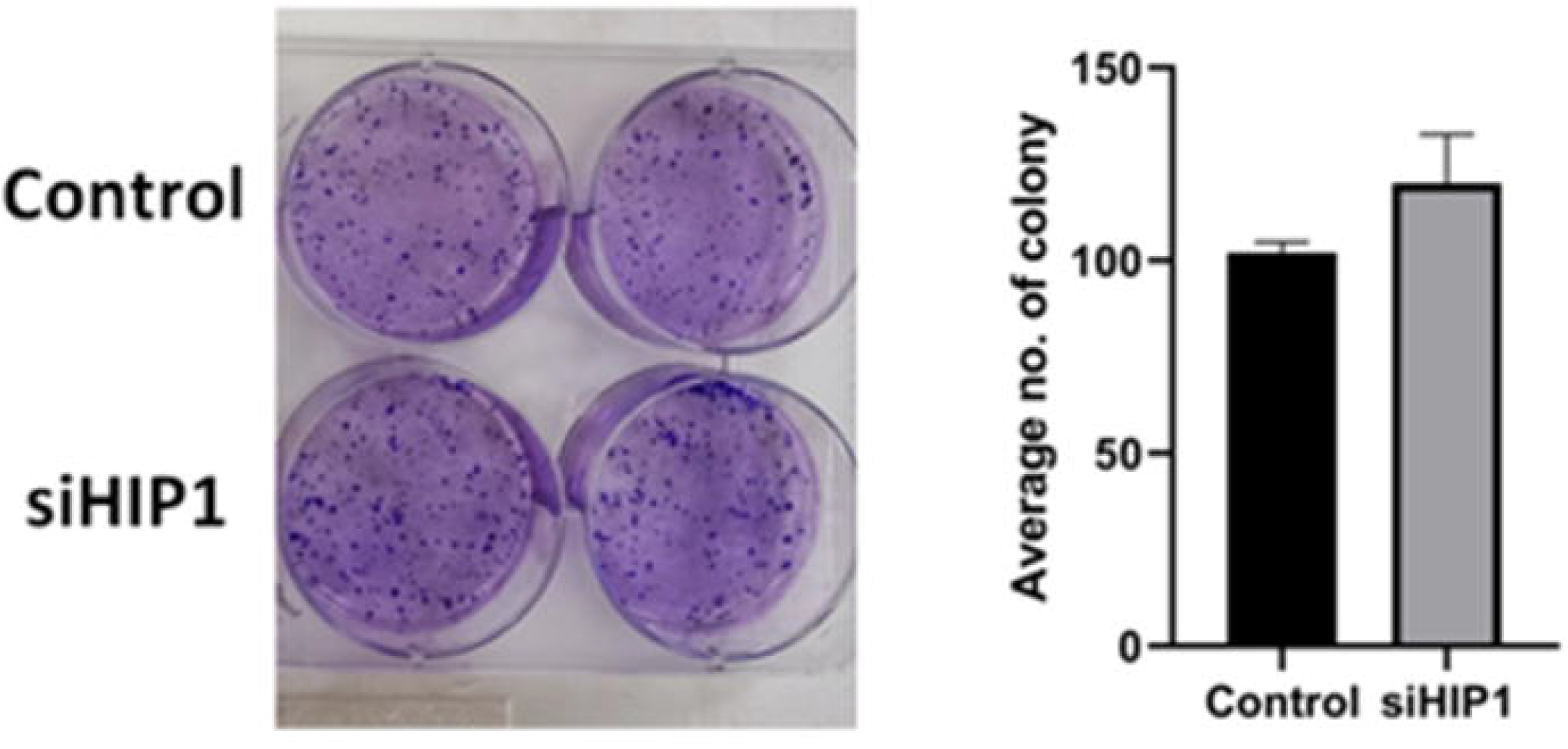
HIP1 siRNA (siHIPl )/scrambled siRNA(Control) was transfected in A549 cells. After 48 hrs cells were seeded into a 6-well plate and allowed to form colonies. After 7 days, cells were fixed, and stained with ciystal violet dye. Colonies were counted and plotted as bar graph (P=0.021).

**Figure 3F.**
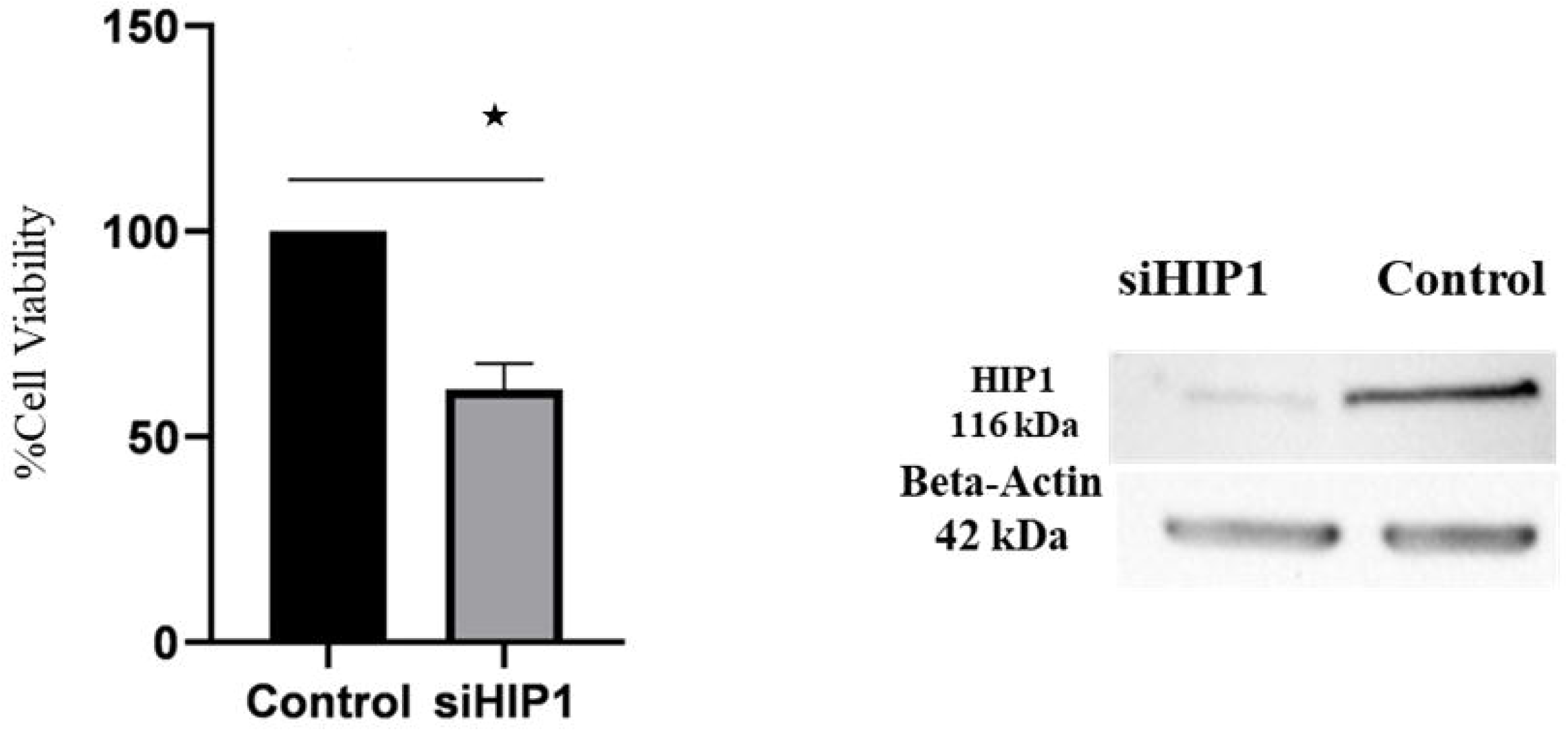
HIP1 siRNA (siHIPl)/scrambled siRNA(Control). After 24 hrs cells were treated with MTT for 3hrs and finally formazan crystals were dissolved by DMSO. Absorbance was taken at 565 nm and percentage cell survival plotted (mitochondrial respiration) as bar graph (p<0.05). Also, HIP1 silencing in the same cells was confirmed using iimnunoblotting.

To investigate HIP1 mediated molecular mechanisms, we compared expression levels of transcripts of markers for cell invasion, proliferation and survival after HIP1 knockdown. Significant induction of proliferation markers, Ki67 (∼5 fold) and PCNA (∼10 fold), was observed. Majority of stemness factors (Oct-4, Nanog, Sox-2, Stat-3) were found reduced, besides Nestin, which was strongly induced by 9 fold. Epithelial marker, E-Cadherin (∼2.4 fold) was found induced, while mesenchymal markers like N-Cadherin (∼2 fold induced) and Vimentin (0.5 fold reduced) were differentially regulated. Most strikingly, a remarkable induction in MMP-9 (∼10 fold) expression, an invasion marker, was observed, which could explain higher invasive abilities of these cells [Fig 3G]. At protein levels, both E-Cadherin and Vimentin expression were significantly induced. However, Zeb-1 was reduced and Beta-catenin was unchanged [Fig 3C].

**Figure 3G:**
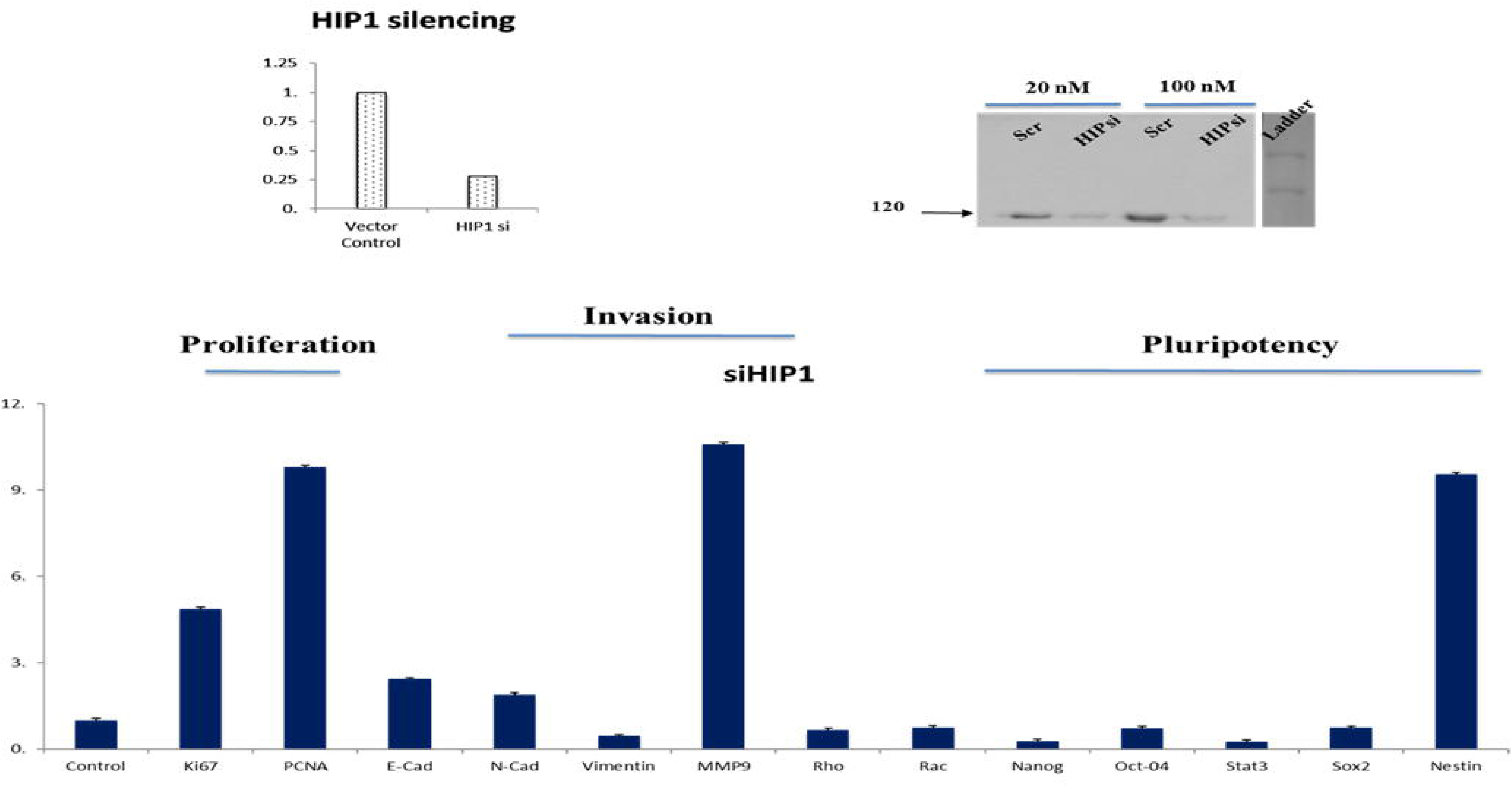
A549 cells were transfected with scrambled siRNA (control) and HIP1 siRNA (siHIPl). After confirmation of HIP 1 knockdown by qRT-PCR and iimnunob lotting, fold change in expression of markers for proliferation (Ki67, PCNA), invasion (E-Cad, N-Cad, Vimentin, MMP9, Rlio, Rac) and pluripotency (Nanog, Oct4, Stat3, Sox2, Nestin) were analyzed by qRT-PCR. Fold change is calculated by 2 ^ddCt^ method and plotted as Fold Change±S.D (Standard Deviation). All the fold changes are significant (p<=0.05))

Additionally, transcript co-expression data for the above genes and HIP1 was explored using cbioportal in Lung Adenocarcinoma patients (OncoSG, Nat Genet 2020) (N=169) [Fig 3H-a, b, c, d]. Here, we observed a significant positive correlation of HIP1 with epithelial markers such as E-Cadherin, Occludin and CD44 [Fig 3H-b] and negative correlation with mesenchymal markers like N-Cadherin (CDH2), SNAIL1 and Twist1 [Fig 3H-c]. A negative correlation was also observed between HIP1 and stemness associated factors, PROM-1 (CD133), SOX-2, MYC [Fig 3H-d]. Proteases critical for invasion, such as MMP-3 and 9 [Fig 3H-c], were also found to be negatively correlated with HIP1 expression. Intriguingly, we also observed a significant positive correlation between a hypoxia-specific transcription factor, EPAS1 (HIF2) and HIP1 [Fig 3H-a].

**Figure 3H-a:**
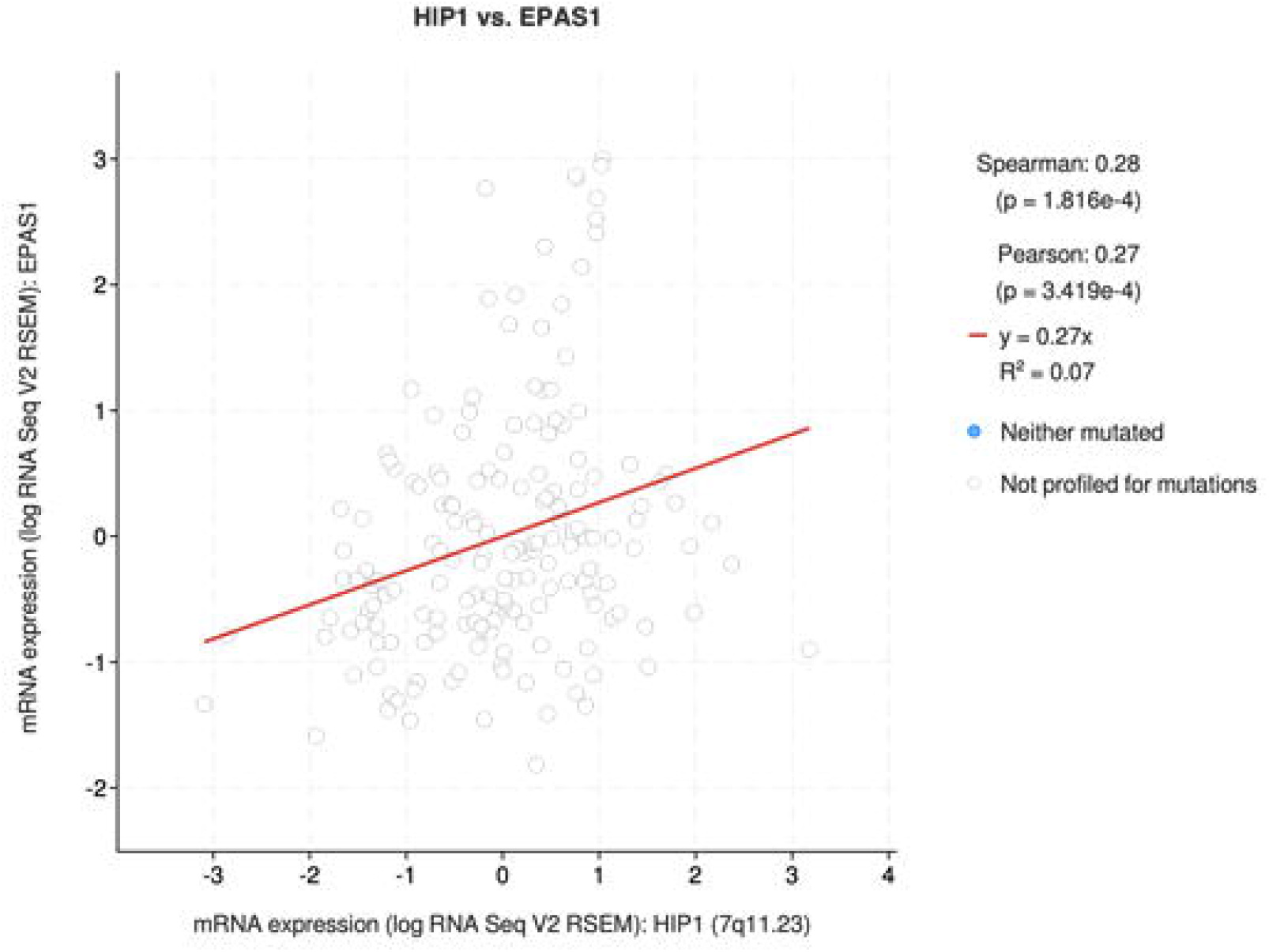
mRNA correlation graph between EPAS1 (HIF2) and HIP1 gene (169 samples) analyzed using cbioportal website.

**Figure 3H-b:**
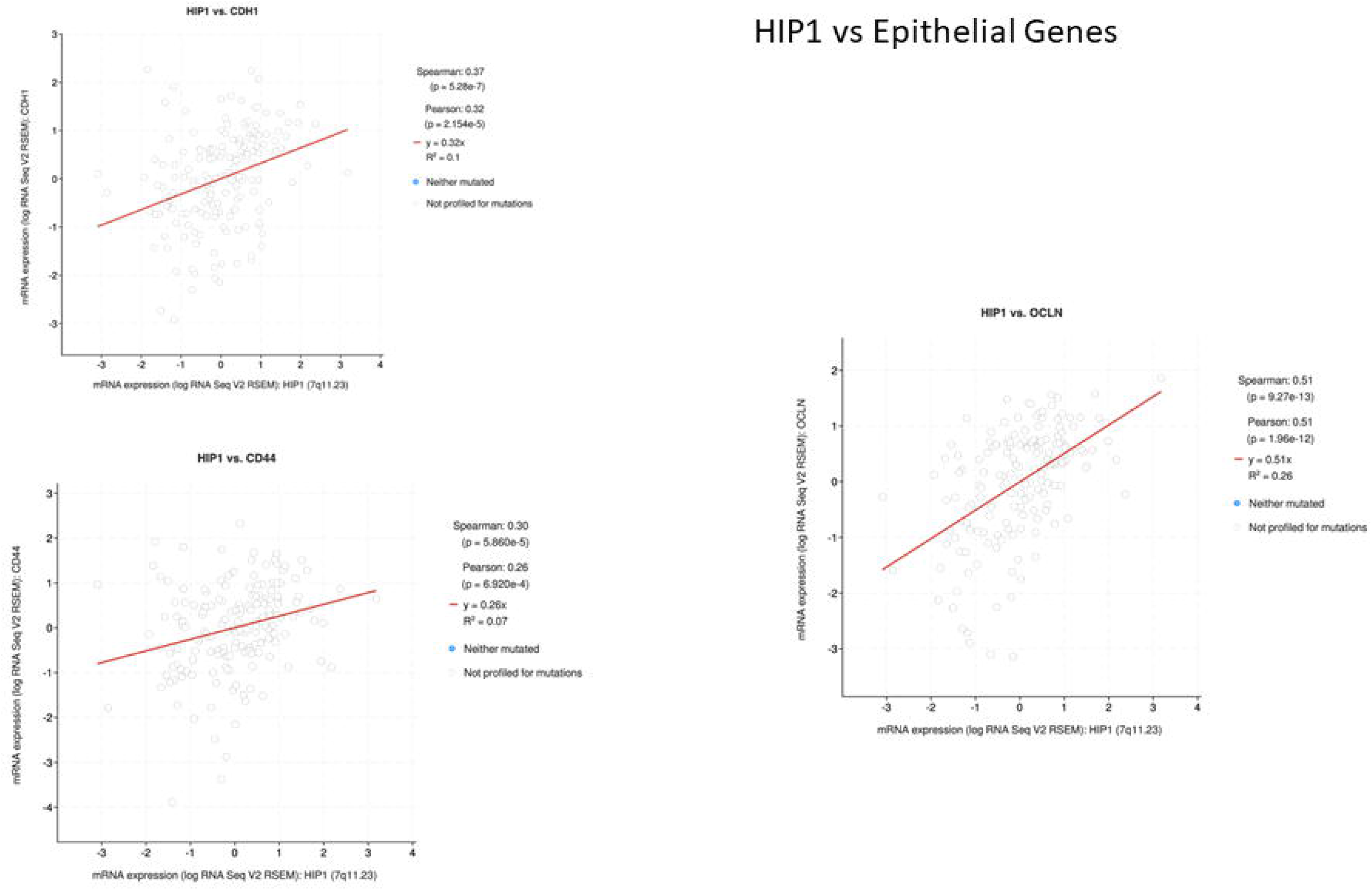
inRNA correlation graph of HIPl with E-Cadlierin, Occludin and CD44 genes. Significant positive correlation was observed between HIP1 and other genes.

**Figure 3H-c:**
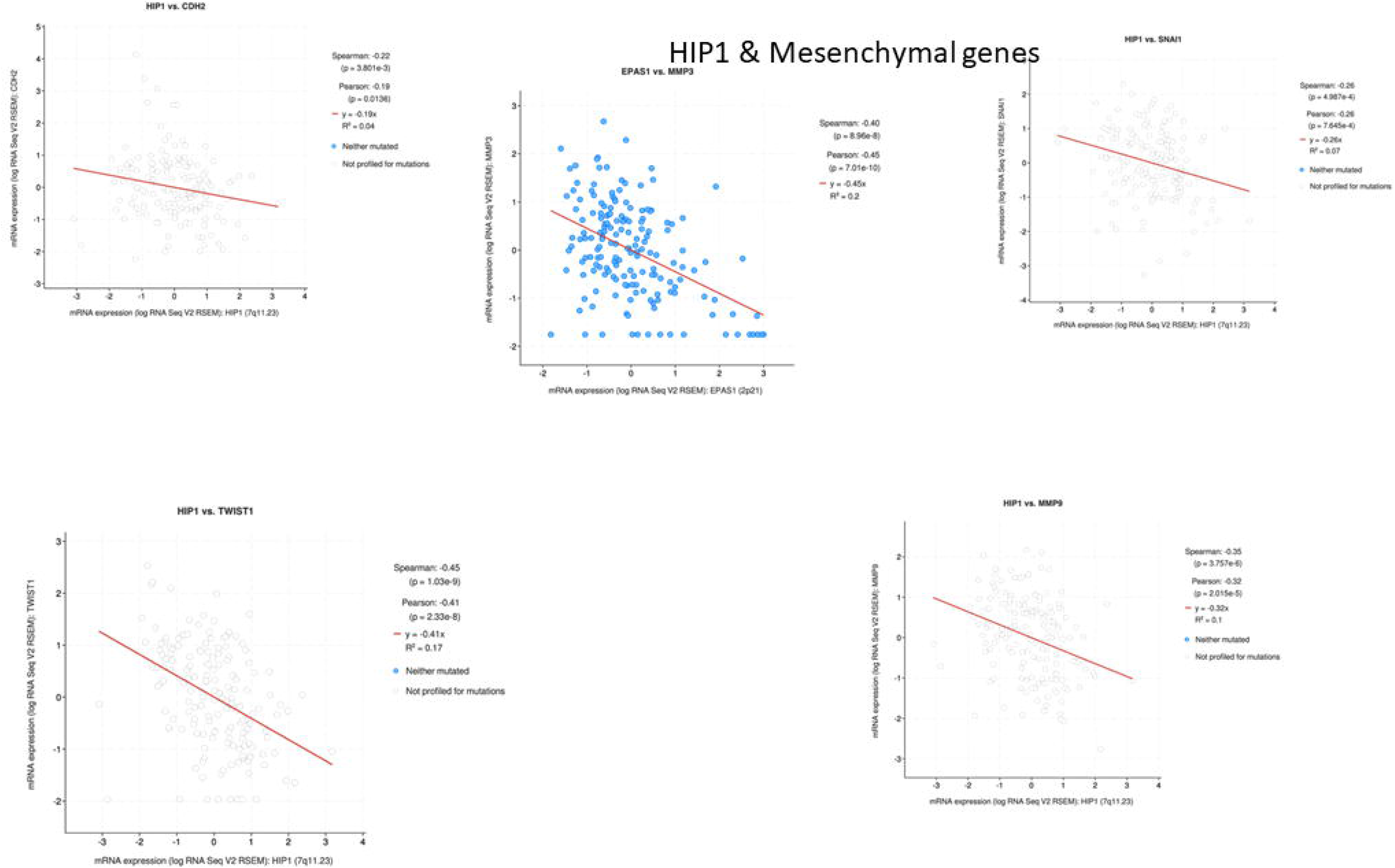
mRNA correlation graph of HIP 1 with mesenchymal markers like N-Cadherin (CDH2). SNAIL1. Twistl and invasion associated proteases like MMP9 and MMP3. Significant negative correlation was observed between HIP 1 and other genes

**Figure 3H-d:**
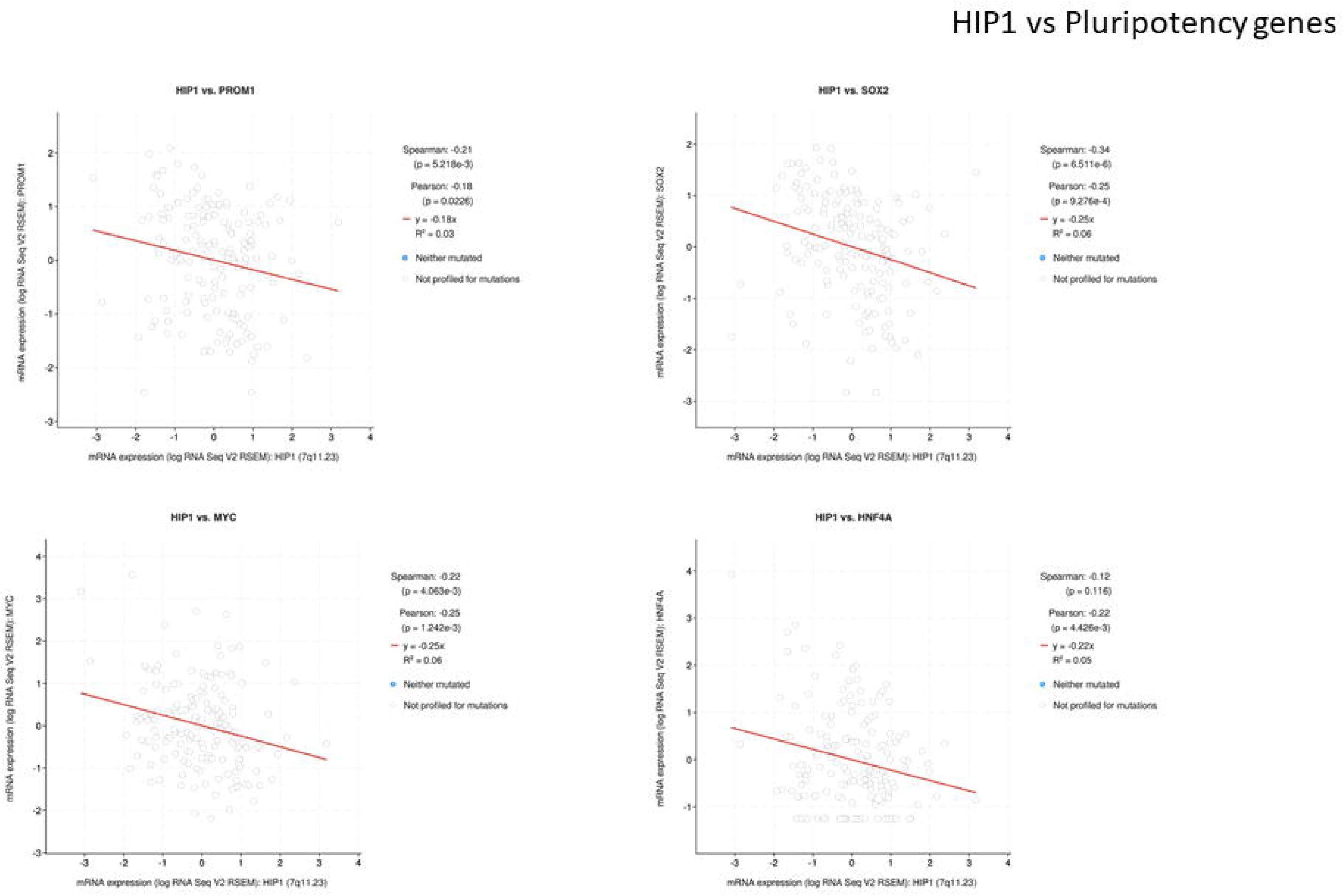
**mRNA correlation graph** of HIP1 with sternness associated factors, PROM-1 (CD133), SOX-2, MYC Significant negative correlation was found between HIP1 and other genes.

### 4. Hypoxic induction of HIP1 expression is mediated by the HIF2 axis

Hypoxic microenvironment, being a cardinal feature of solid tumours, is involved in invasiveness, drug resistance and stemness. As a strong correlation was observed between HIF2 and HIP1 transcript expression using TCGA data and we also observed predicted induction of NFE2L2 (NRF2) mediated pathways from our proteomics data, we further explored the possibility of regulation of HIP1 expression in hypoxic conditions.

Expression of HIP1 was studied in lung cancer cells under hypoxic conditions (1% oxygen) at various time intervals. We observed a marked increase in HIP1 transcript and protein levels under hypoxic conditions (Fig 4A). Additionally, HIP1 expression was also found to be differentially regulated by hypoxia in other cancer types (data not shown). To further investigate if hypoxia inducible factors, HIF1 or HIF2, mediate hypoxic induction of HIP1, we overexpressed them using transient transfections of their mutant stabilising constructs. Induction of VEGF transcript levels was observed in both HIF1 and HIF2 overexpressed cells, confirming activation of their downstream targets. However, expression of HIP1 transcript (1.7 fold) and protein (1.4 fold) were induced specifically only in HIF2 overexpressed cells. Thus, HIF2 induces HIP1 in chronic hypoxic conditions. This could be a possible feedback mechanism employed by hypoxic tumour cells to enable their cell survival under these conditions.

**Figure 4A:**
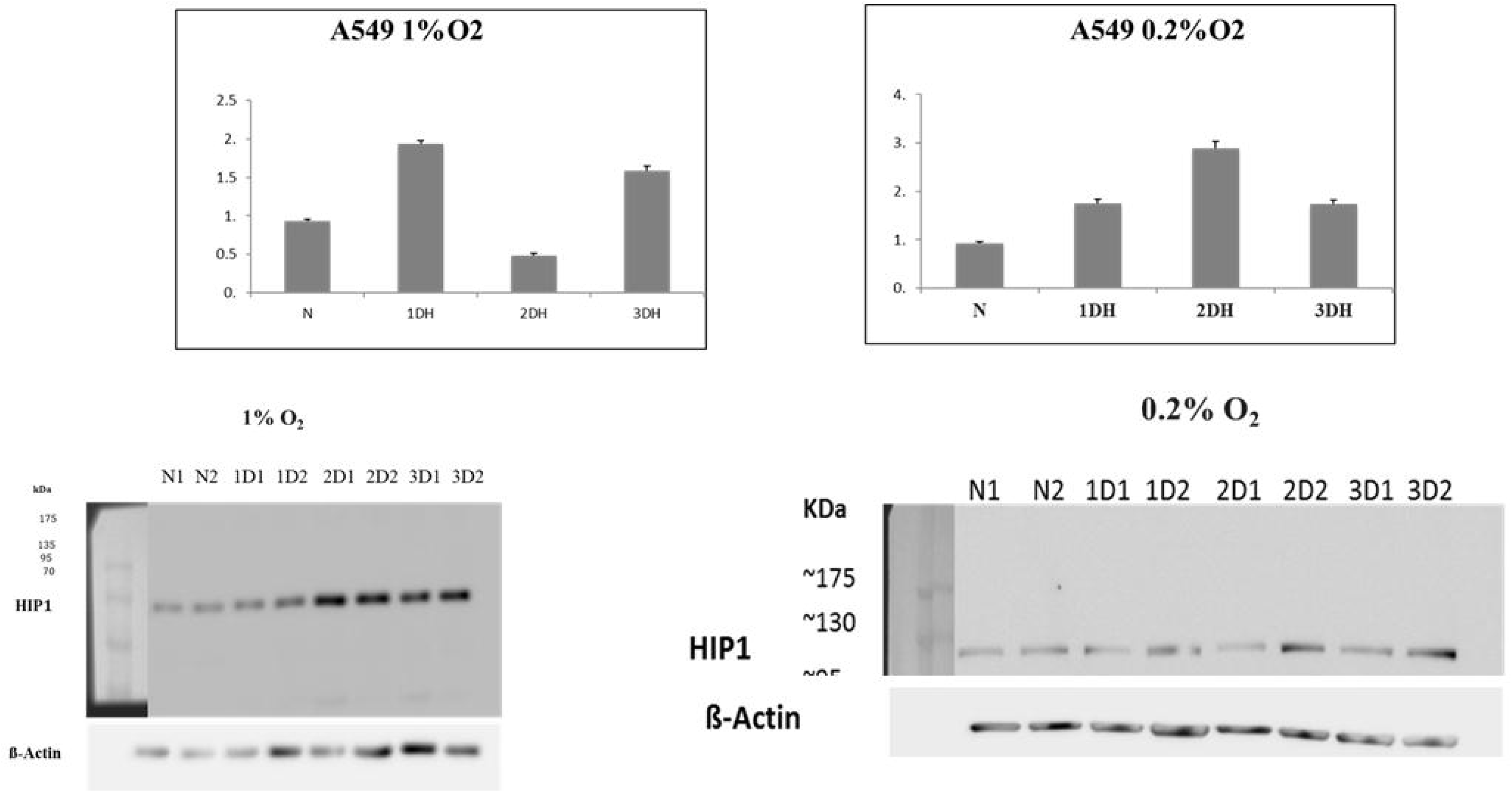
A549 cells were cultured in different hypoxic conditions (1%O_2_ and 0.2%O_2_) for 24 hours (1DH), 48 horns (2DH) and 72 hours (3DH). The differential expression of HIP1 due to hypoxia was analyzed by qRT-PCR and immunoblotting by using beta actin as controls. Fold change is calculated by 2 ^ddCt^ method and plotted as Fold Change±S.D (Standard Deviation).

**Figure 4B:**
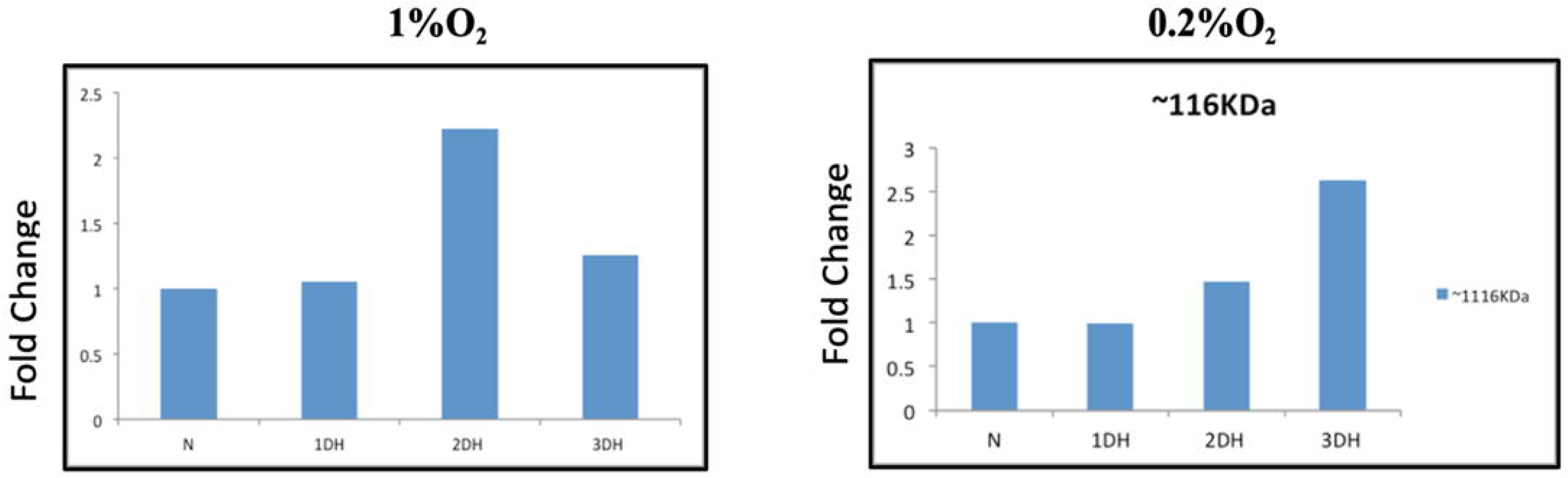
Densitometric analyses of the bands obtained from immunoblotting plotted using normalized values as fold change compared to normoxia.

**Figure 4C:**
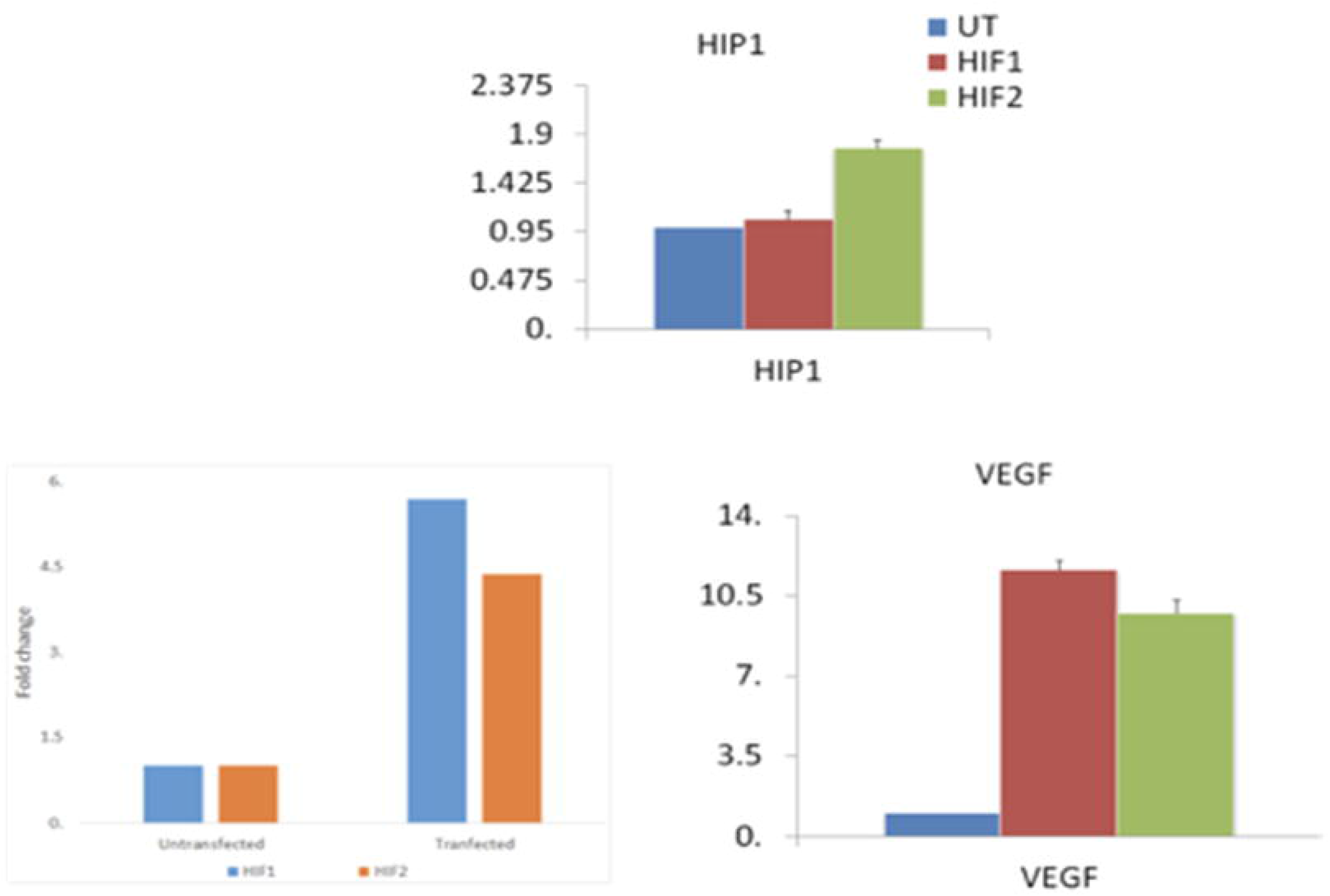
A549 cells were transiently overexpressed with mutant HIF-1 and mutant HIF-2 plasmids The effect of HIF on HIP1 expression was analyzed by qRT-PCR compared to the untransfected (UT) control. Induction of HIF was confirmed by analyzing induction of VEGF with respect to UT.

**Figure 4D:**
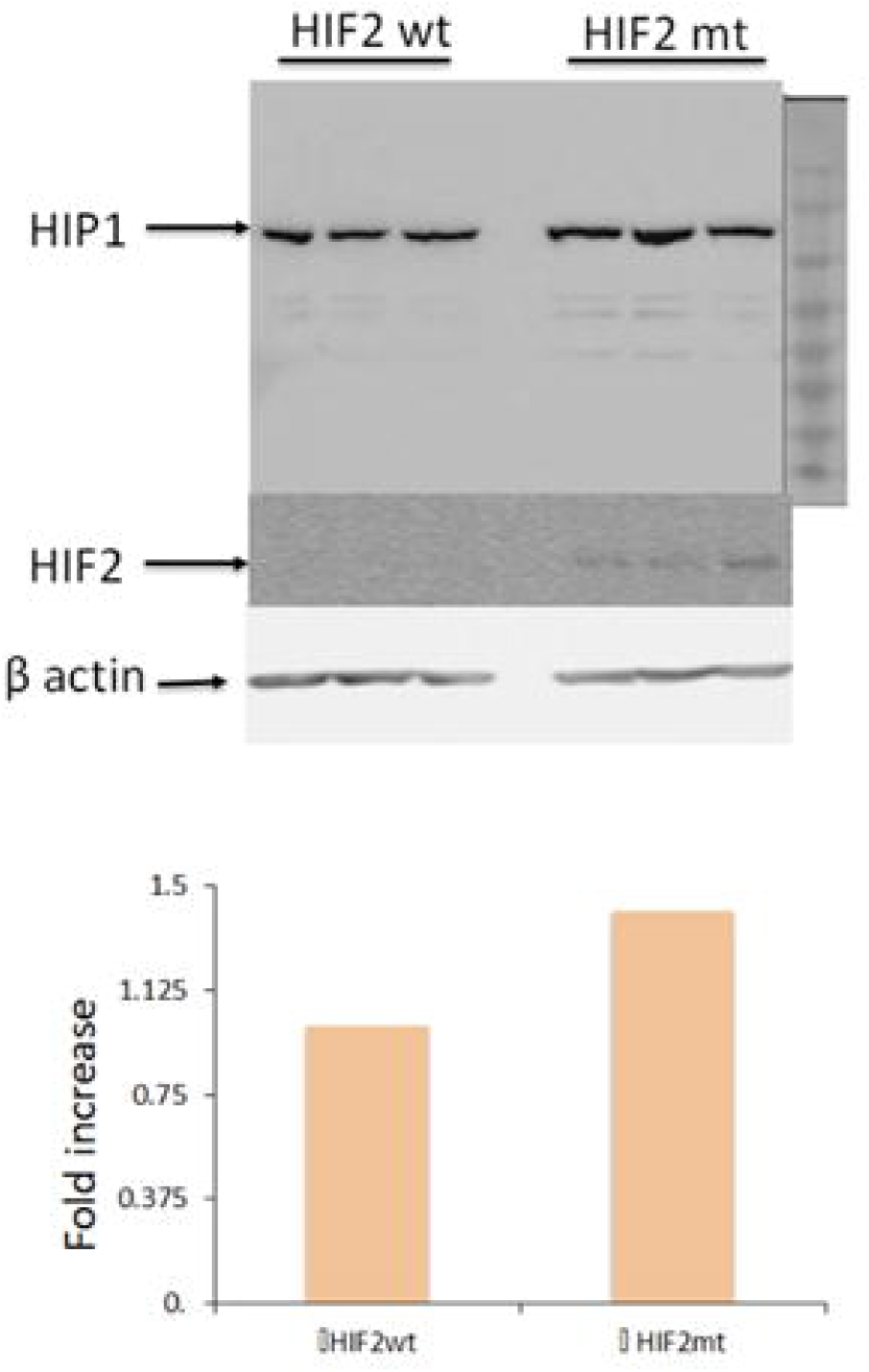
A549 cells were transiently overexpressed with wtHIF-2 and mtHIF-2. The effect of wtHIF-2 and mtHIF-2 on HIP1 expression was analyzed by immunoblotting. The difference in HIP1 expression was determined as fold change with respect to beta-actin by densitometry.

## DISCUSSION

Lung adenocarcinoma presents with high mortality rates and is usually diagnosed at advanced stages. In order to improve patient prognosis, there is an unmet need for early markers and novel targets. *Kras* mutant NSCLC is the most common type and inspite of MEK and EGFR targeted therapies, relapse occurs because of feedback activation of other RTKs [15]. Our study provides novel insights on regulation of an endocytic protein, HIP1, in an *in vitro Kras* mutant lung adenocarcinoma model. Besides identifying Hypoxia Inducible Factor 2 (HIF2) as an upstream regulator, our proteomics data implies that HIP1 regulates various cellular processes, majorly, metabolism and RNA transport. Patient data analysed from TCGA and functional assays for migration and invasion also corroborate earlier findings that HIP1 expression is reduced at advanced stages of lung cancers and has an anti-metastatic role.

Analysis of TCGA data demonstrates that HIP1 expression is reduced in various cancer types including NSCLC compared to matched or unmatched normal tissues. RNA expression data from patient samples reveals that HIP1 expression is directly correlated to overall survival in NSCLC subtypes. Metastatic tumor tissues had least HIP1 expression and significantly reduced overall survival. This corroborates earlier findings by Hsu et al. (2016) demonstrating reduced HIP1 protein levels in tissue samples from advanced stages of lung adenocarcinoma [10]. Also, transcript co-expression data from TCGA reveals a positive correlation of HIP1 with epithelial markers, such as E-Cadherin, Occludin and CD44, and negative association with mesenchymal markers like N-Cadherin (CDH2), SNAIL1 and Twist1. Robust transcript induction of MMP9, known matrix protease for metastasis [16] and Nestin, known cancer stem cell marker [17], indicates both these processes to be activated by HIP1 silencing. Also, our functional migration and invasion assays confirm HIP1’s anti-metastatic role. Furthermore, induction of FGFR1 and 2 (1.5fold) in the proteomics data of HIP1 silenced cells, confirms their mesenchymal characteristics as demonstrated by Kitai et al. [18]. In fact, HIP1 regulated cell survival enabling anchorage independent growth in soft agar assay, also suggest that HIP1 may play a role in anoikis resistance and stemness. Contrary to the functional assays and supporting data, we have not observed the canonical regulation of EMT markers using immunoblotting. Significant induction of protein levels of both E-Cadherin and Vimentin together imply activation of non-classical pathways of EMT, such as collective migration, which have higher metastatic abilities [19]. This aspect needs to be explored in further details.

Unlabeled quantitative mass spectrometry analysis was performed to identify globally deregulated proteome after silencing HIP1 expression. Metabolic pathways were found to be most perturbed and consequently, a significant Succinate Dehydrogenase B (SDHB) depletion (80%) was validated. As SDHB is a key enzyme for mitochondrial respiration, its depletion may shift metabolism from aerobic respiration to glycolysis. Reduced MTT readings indicative of redox potential also imply this. Similarly reduced SDHB expression is reported in advanced stages of Hepatocellular carcinoma [20, 21]. Impaired activity of SDHB may contribute to oxidative stress because of reduced superoxide anion scavenging ability. SDHB is also shown to function as a tumour suppressor, its inhibition leading to EMT and enhanced proliferation in ovarian cancer [22]. SDHB deficient cells have impaired mitochondrial respiration and increased glycolysis [23]. We have also similarly found activated pathways leading to increased PFKP and LDHA in our proteomics data (Fig 2E). Regulatory network analysis of HIP1 silenced deregulated proteome data presented with MYC, a well-studied proto-oncogene, upstream to NFE2L2/NRF2 activation. NRF2 is a transcription factor involved in cellular resistance to oxidative damage [24]. It activates the genes key to redox homeostasis, drug metabolism and excretion, energy metabolism, survival, proliferation and mitochondrial physiology [25]. KEAP1 (Kelch-like ECH-associated protein 1), which is also a redox sensitive protein, is a repressor of NFE2L2. Under physiological homeostasis, NRF2 is bound by KEAP1, which causes its proteasomal degradation. KEAP1 is mutated in A549 cells and is less expressed, keeping NRF2 pathways usually activated. Sustained activation of NRF2 drives resistance to chemotherapeutic drugs [26] and also activates the kynureninase pathway which leads to tumour promotive and immune suppressive effects in lung cancers [27]. Hence further studies are required to uncover the significance of MYC and NRF2 activation in response to HIP1 depletion and explore if these could be important targets at advanced stages.

All solid tumours have hypoxic conditions, and cancer cells undergo various adaptations such as increased glycolysis [28], angiogenesis, anoikis resistance [29] and stemness [30] to survive and proliferate in this microenvironment. Our study is the first to demonstrate that HIP1 expression is regulated by hypoxic stress conditions, mediated specifically via HIF2, as the upstream regulator. Also, hypoxic conditions render the cells more invasive thereby accelerating disease progression. The key hypoxia inducible transcription factors, HIF1 and HIF2, have distinctive and sometimes opposing roles in the same cell, acting through independent target genes and binding to contrasting regulators [31]. Contrary to studies demonstrating HIF2 overexpression as a negative prognostic indicator in NSCLC, it functions as a tumour suppressor in *Kras* dependent tumours by suppressing Akt pathway through induction of HIN-1 [32]. Here we demonstrate that HIP1 is induced under hypoxic conditions and the HIF2 axis mediates this response independent of HIF1. HIP1 is also shown to inhibit the Akt-beta catenin pathway, thereby acting as a metastatic suppressor in NSCLC [10]. Thus, our findings corroborate the anti-metastatic role of HIP1 and imply that HIF2 may suppress Akt activity by regulating both HIP1 and HIN-1, thereby acting as a potent tumour suppressor in *Kras* driven NSCLC cells. Additional studies in the context of a hypoxic microenvironment and validation of identified targets like C-MYC and NRF2 will carry significant theragnostic implications.

