## Supplementary figures for "Huntingtin Interacting Protein-1 expression is regulated via HIF2 axis in Lung Adenocarcinoma"

Supplementary Figure 1

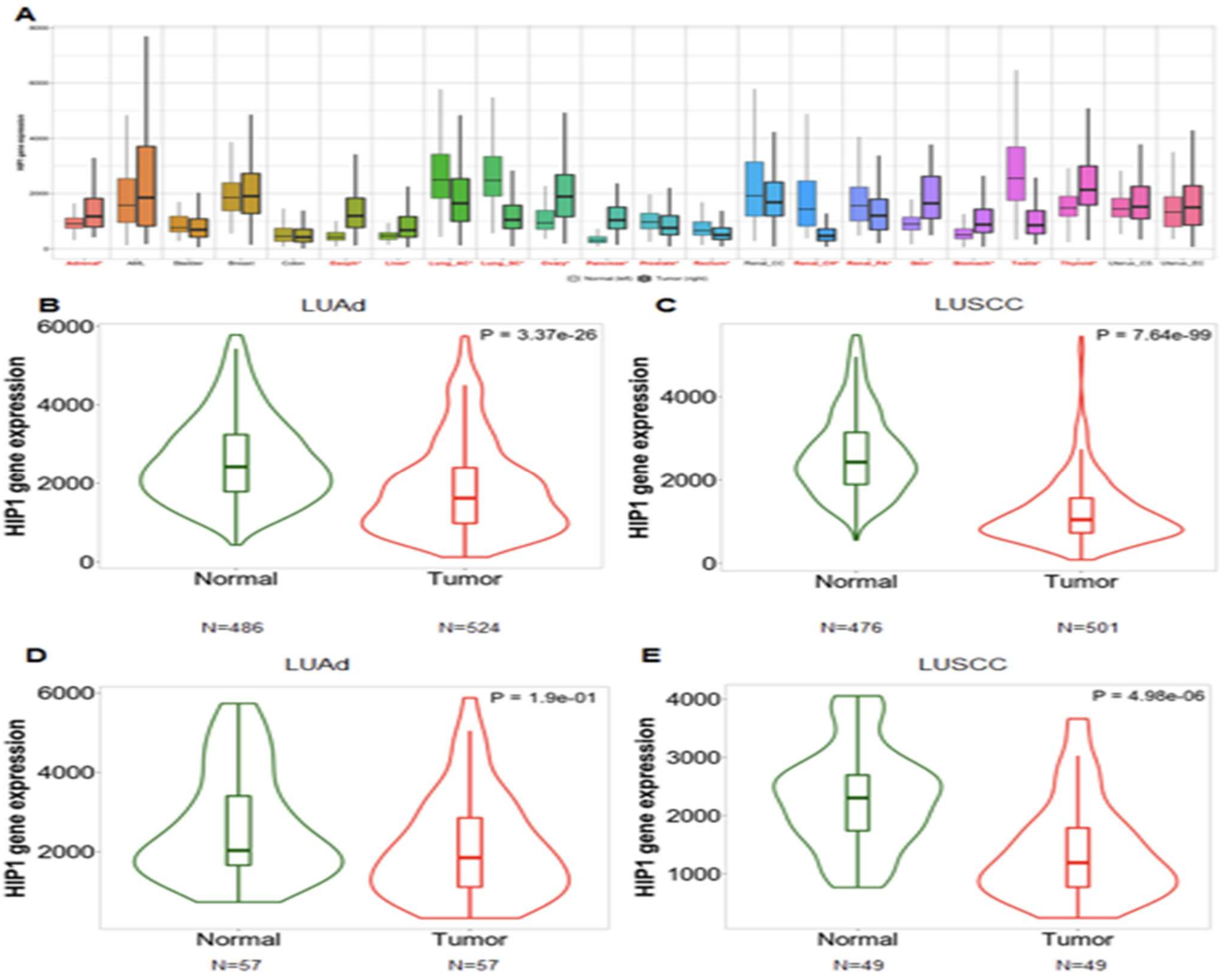


**Supl Figure 1**: HIP1 expression in lung cancer. (A) HIP1 gene expression in tumor cells compared to normal tissues among all cancer types. (B,C) RNA-seq data for lung adenocarcinoma (LUAd) and lung squamous cell carcinoma (LUSCC) for HIP1 expression in tumor tissues compared to normal tissues including normal from non-cancerous patients. (D,E) RNA-seq data for HIP1 expression in paired tumor-normal tissues of LUAd and LUSCC. Significant differences by Mann-Whitney U test are marked in red*.

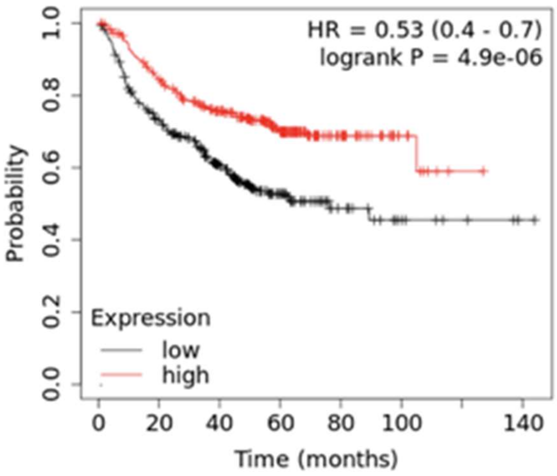


**Supl Figure 2**: Kaplan-Meier survival curves of lung cancer patients with decreased expression of HIP1 showed poor progression free survival compared to those with increased HIP1 expression.

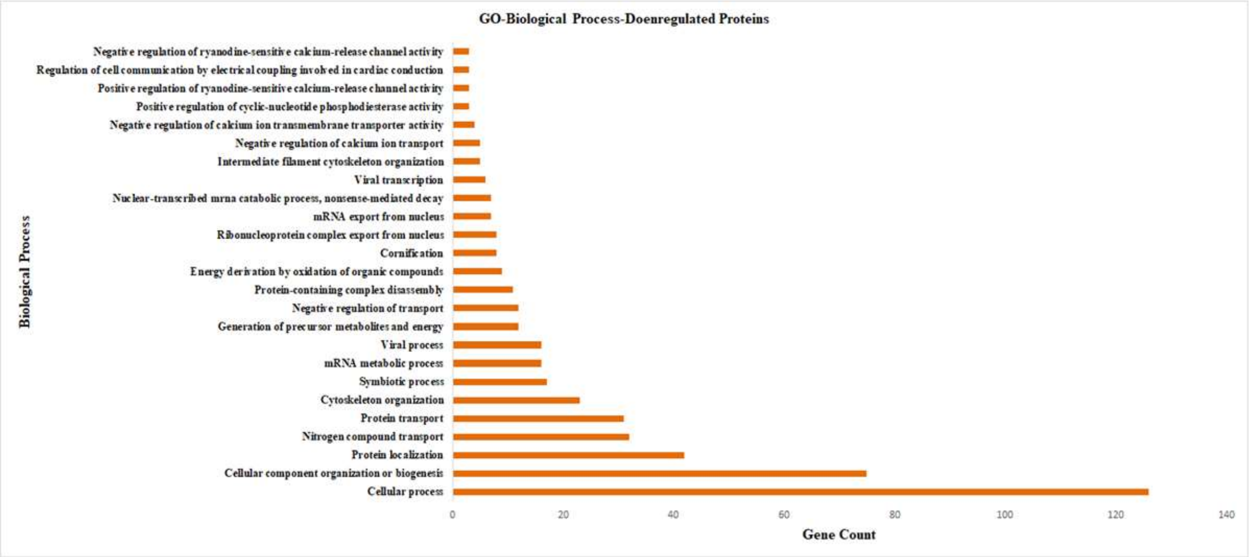


**Supl Figure 3**. The downregulated proteins found after proteomic profiling of HIP1 silenced cells were analysed for their involved biological processes using STRING software. Major downregulated biological processes thus found are shown as a bar diagram. Y axis denotes the gene count and X axis denotes the biological process.

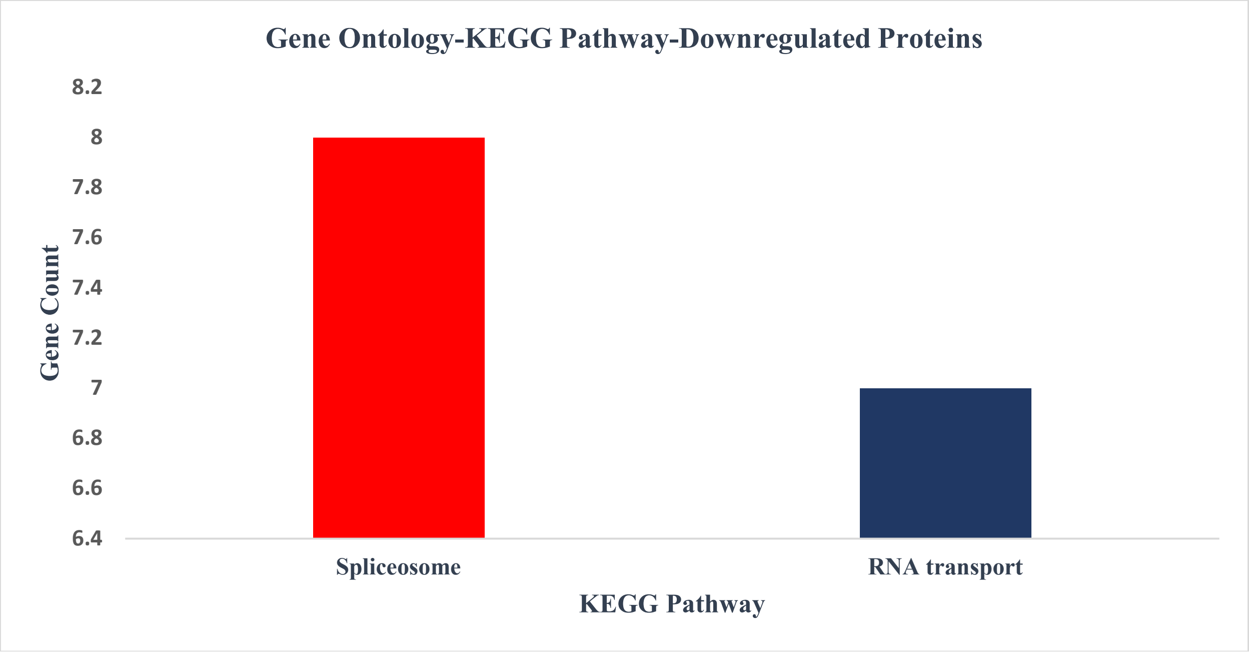


**Supl Figure 4.** The downregulated proteins found after proteomic profiling of HIP1 silenced cells were analysed for their involved KEGG pathways using STRING software. Major downregulated KEGG pathways thus found are shown as a bar diagram. Y axis denotes the gene count and X axis denotes the KEGG pathways.

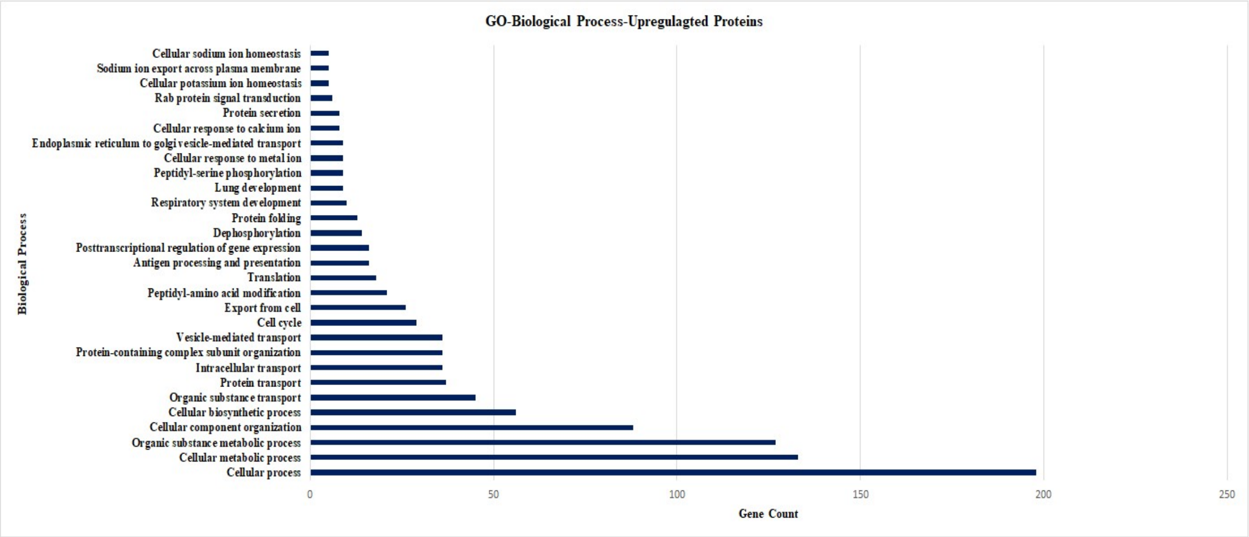


**Supl Fig 5**. The upregulated proteins found after proteomic profiling of HIP1 silenced cells were analysed for their involved Biological processes using STRING software. Major upregulated biological processes thus found are shown as a bar diagram. Y axis denotes the biological process and X axis denotes the gene count.

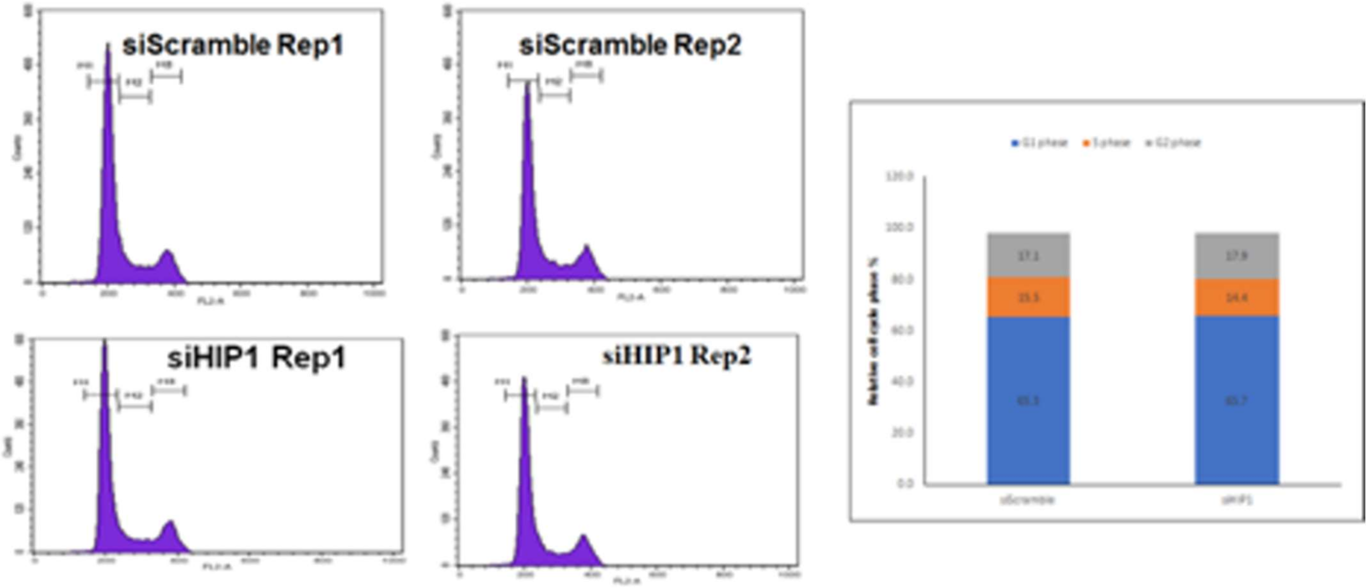


**Supl Fig 6.** HIP1 siRNA (siHIP1)/scrambled siRNA(Control) was transfected in A549 cells. After 48 hrs cells were harvested and fixed in 70% ethanol, treated with ribonuclease and stained with propidium iodide (PI). Cells were then analyzed by flow cytometry and % change in G1 (blue color), S (orange color) and M phase (grey color) plotted as stacked bar graph.

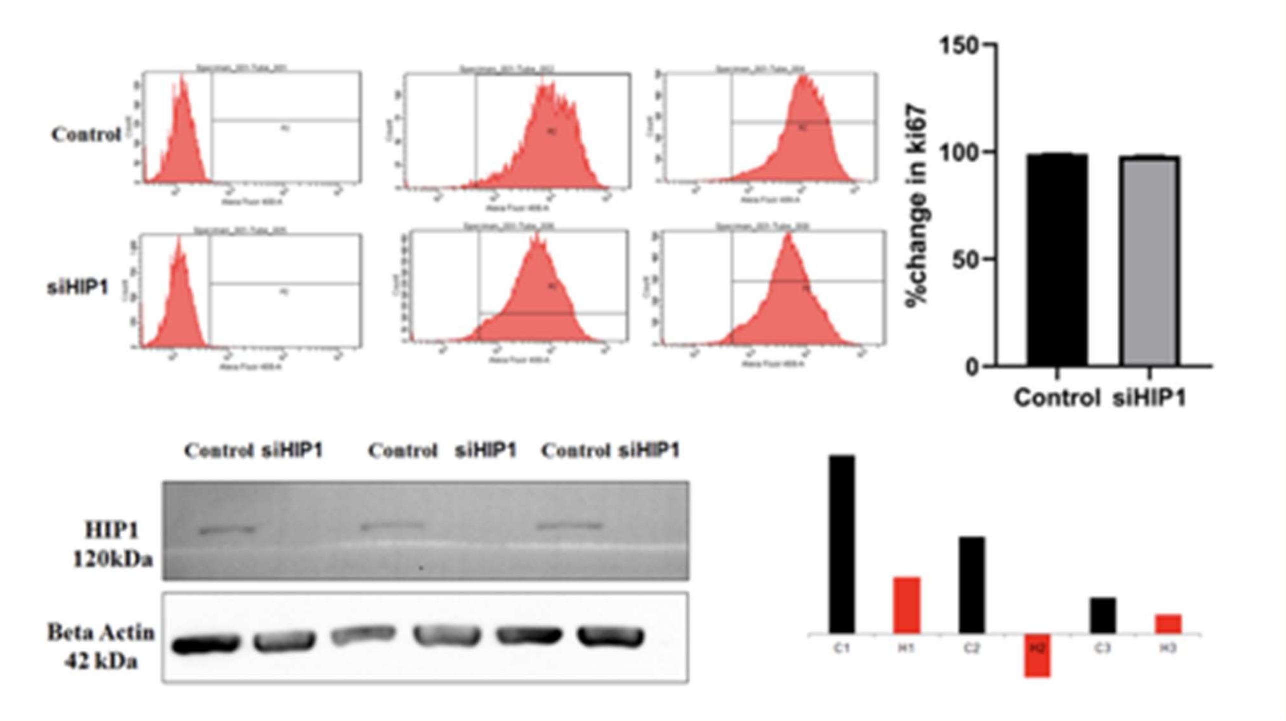


**Supl Fig 7**. HIP1 siRNA (siHIP1)/scrambled siRNA(Control) was transfected in A549 cells. After 48 hrs cells were harvested and permeabilized by 0.2% triton x, stained with Ki67 antibody. After this cells were washed and resuspended in staining buffer and analyzed by FACS. % change in ki67 level was plotted as bar graph. Also, HIP1 silencing in the same cells was confirmed using immunoblotting.
